# RNA splicing analysis using heterogeneous and large RNA-seq datasets

**DOI:** 10.1101/2021.11.03.467086

**Authors:** Jorge Vaquero-Garcia, Joseph K. Aicher, Paul Jewell, Matthew R. Gazzara, Caleb M. Radens, Anupama Jha, Christopher J. Green, Scott S. Norton, Nicholas F. Lahens, Gregory R. Grant, Yoseph Barash

## Abstract

The ubiquity of RNA-seq has led to many methods that use RNA-seq data to analyze variations in RNA splicing. However, available methods are not well suited for handling heterogeneous and large datasets. Such datasets scale to thousands of samples across dozens of experimental conditions, exhibit increased variability compared to biological replicates, and involve thousands of unannotated splice variants resulting in increased transcriptome complexity. We describe here a suite of algorithms and tools implemented in the MAJIQ v2 package to address challenges in detection, quantification, and visualization of splicing variations from such datasets. Using both large scale synthetic data and GTEx v8 as benchmark datasets, we demonstrate that the approaches in MAJIQ v2 outperform existing methods. We then apply MAJIQ v2 package to analyze differential splicing across 2,335 samples from 13 brain subregions, demonstrating its ability to offer new insights into brain subregion-specific splicing regulation.

## Introduction

The usage of RNA sequencing (RNA-seq) has become ubiquitous in biomedical research. While some studies utilize RNA-seq only to investigate the overall expression level of genes, an increasing number of studies analyze changes in the relative abundance of gene isoforms. Changes in gene isoforms can occur through multiple mechanisms, including alternative promoter usage, alternative polyadenylation, and alternative splicing (AS). The production of different gene isoforms can in turn lead to diverse functional consequences, including changes to the translated protein domains, to degradation rates, and to localization. Previous studies showed that the majority of human genes are alternatively spliced with over a third of them shown to change their major isoform across 16 human tissues (*1*). These observations, combined with the association of splicing defects with both monogenic and complex disease, serve to motivate the study of splicing variations across diverse experimental conditions. Consequently, independent labs as well as large consortia produce vast amounts of RNA-seq data. Datasets may involve anywhere from just a few to many thousands of samples each, and are typically heterogeneous as they often do not represent biological or technical replicates. The consequent increased splicing variability, illustrated in Fig 1A,B, can be the result of a multitude of factors, both experimental (e.g. difference in sequencing machine), and biological (e.g. gender, age). While some confounding factors may be corrected with appropriate methods (*2*), fully removing the observed variability in such data is unlikely and may also over-constrain the data, thus leading to a loss of true biological signal. Thus, there is a general need for methods that can effectively detect, quantify, and visualize splicing variations from large and heterogeneous RNA-seq datasets.

**Fig. 1:**
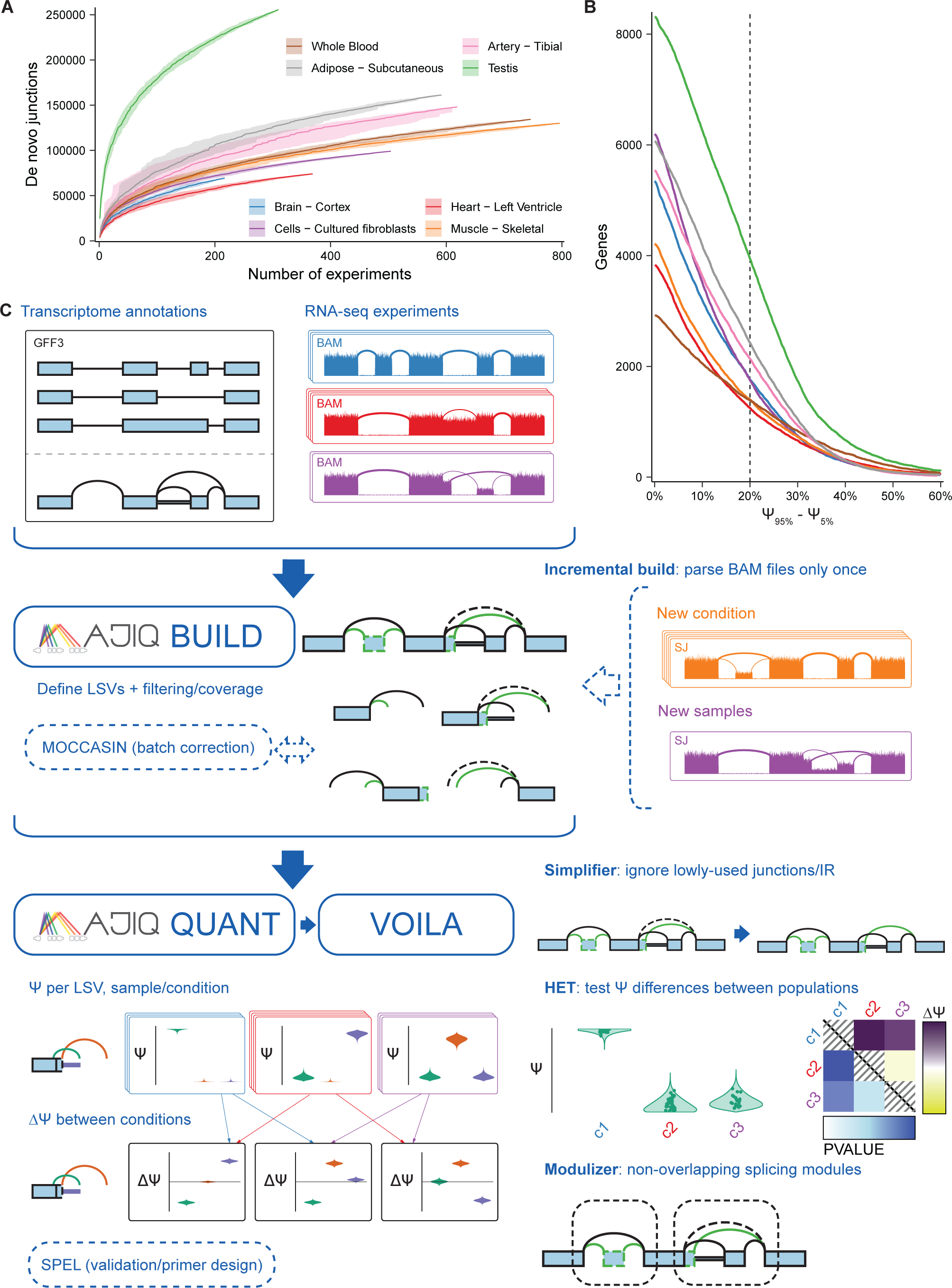
MAJIQ efficiently and accurately models, quantifies, and visualizes RNA splicing from large and complex RNA-seq datasets. **(A)** The number of identified distinct unannotated *de novo* junctions increases with larger subsets of different tissues from GTEx. Lines show the median over 30 randomly selected permutations over experiments in each subset, confidence bands show the 5th to 95th percentiles over permutations of samples per tissue. **(B)** The number of genes with at least one junction where the difference between the 95th percentile and 5th percentile of PSI exceeds a given value for different tissues from GTEx (same tissues/colors as in Fig. 1A). Dashed vertical line indicates how many genes have a difference in PSI exceeding 20%. **(C)** MAJIQ combines annotated transcript databases and coverage from input RNA-seq experiments to build a model of each gene as a collection of exons connected by annotated and *de novo* junctions and retained introns (splicegraph). Junctions and retained introns sharing the same source or target exon form local splicing variations (LSVs). MAJIQ quantifies the relative inclusion of junctions and retained introns in each LSV in terms of percent spliced in (PSI, Ψ) and provides VOILA to make interactive visualizations of splicing quantifications with respect to each gene’s splicegraph and LSV structures. MAJIQ v2 introduces an incremental build, which allows RNA-seq coverage to be read from BAM files only once to a coverage file (SJ), accelerating subsequent builds with different experiments. MAJIQ v2 introduces a simplifier, which can be used to reduce splicegraph/LSV complexity by ignoring lowly used junctions and retained introns. MAJIQ v2 introduces a new mode for quantification, HET, which compares PSI differences between populations of independent RNA-seq experiments and accounts for variable uncertainty per experiment. MAJIQ v2 introduces the modulizer, which allows performing analysis relative to non-overlapping splicing modules rather than LSVs.

Broadly, the quantification of changes in gene isoform usage can be divided between methods that aim to quantify whole isoforms and those that quantify more local AS “events” within a gene. While quantifying all gene isoforms accurately across diverse conditions can be regarded as the grand challenge of transcriptomics, achieving this goal remains open due to several limiting factors. In the case of long reads technology, these factors include high error rate and high costs which do not allow researchers to capture enough reads from all isoforms. In the case of the more commonly used short reads technology, these limiting factors include the sparsity of reads, their positional bias, and the fact that reads usually cannot be assigned to a unique isoform. In addition, the composition of isoforms in a sample is typically unknown, requiring further inference of the existing isoforms or making simplifying assumptions such as a known transcriptome. These issues have led many researchers to focus on local AS “events” which can be more easily and accurately quantified from RNA-seq. AS events are quantified in terms of percent spliced in (PSI, denoted by Ψ), which is the relative ratio of isoforms including a specific splicing junction or retained intron. Traditionally, AS events have been studied only for a restricted set of the most common “types” (e.g. cassette exons). In a previous study, we extended this set of AS event types using the formulation of local splicing variations (LSVs) and introduced MAJIQ as a software package for studying such LSVs. LSVs, which can be defined as splits in a gene splicegraph coming into or from a reference exon, allow researchers to capture not only previously defined AS types but also much more complex variations involving more than two alternative junctions (see examples in Figure 1C for illustration). Furthermore, the LSV formulation, and similar definitions of local AS events suggested in subsequent works, also help incorporate and quantify unannotated (*de novo*) splice junctions. Previous work comparing splicing across mouse tissues has shown that accounting for complex and *de novo* variations results in over 30% increase of detected differentially spliced events while maintaining the same level of reproducibility (*3*). Importantly, capturing such unannotated splice variations is of particular importance for the study of disease such as cancer and neurodegeneration which often involve aberrant splicing.

Despite previous demonstrations of MAJIQ’s utility for analyzing AS (*3, 4*), we found it as well as many other commonly used methods for AS events quantification not to be well-suited for handling heterogeneous and large RNA-seq datasets. Such datasets pose several algorithmic, computational, and visualization challenges. First, the assumption of a shared PSI per LSV junction in a group, used by methods such as MAJIQ and LeafCutter, is violated in such data even when handling only a small dataset with few samples, leading to a potential increase in false positives and loss of power. Second, algorithms need to not only scale to thousands of samples efficiently but also to allow incrementally adding new samples as more data is acquired, and to support multiple group comparisons (e.g. multiple tissue comparisons across GTEx). Third, the increased complexity of the data requires efficient representation. Such efficient representation would allow users to capture the many unannotated splicing variations in the data, while at the same time simplifying its representation and quantification. Such simplification will allow to filter lowly used splice junctions while also detecting possibly new sub-types of significant variations. Finally, efficient and user-friendly visualization is required to probe possibly multiple sample groups as well as individual samples.

To address the above challenges, we developed an array of tools and algorithms included in the MAJIQ v2 package. These include nonparametric statistical tests for differential splicing (MAJIQ HET), an incremental splicegraph builder, a new algorithm for quantifying intron retention, a method to detect high-confidence negative (non-changing) splicing events, and an algorithm to parse all LSVs across genes into modules which can then be classified into subtypes (Modulizer). These algorithms and tools are coupled with a new visualization package (VOILA v2) which allows users to compare multiple sample groups, simplify splicegraphs, and probe individual data points (e.g. LSV in an individual sample) while representing hundreds or thousands of samples. In addition, to support reproducibility, we develop a package for comparative evaluation of different methods for RNA splicing analysis and use it to demonstrate that the new version of MAJIQ compares favorably with the current state of the art using both synthetic (simulated) and real (GTEx) data. Finally, we apply the MAJIQ v2 toolset to 2,335 RNA-seq samples from 374 donors across 13 brain subregions. We use VOILA v2 to visualize the result and highlight several key findings in brain subregions specific variations in cerebellar tissue groups compared to the remaining brain regions.

## Results

### The MAJIQ v2 splicing analysis pipeline

To support RNA splicing analysis using large RNA-seq datasets we implemented the set of tools and algorithms illustrated in Figure 1C. In the first step, the MAJIQ builder combines transcript annotations and coverage from aligned RNA-seq experiments in order to build an updated splicegraph for each gene which includes *de novo* (unannotated) elements such as junctions, retained introns, and exons). Several user-defined filters can be applied at this stage to exclude junctions or retained introns which have low coverage or are not detected in enough samples in user-defined sample groups. Notably, per-experiment coverage is saved separately so that it can be used in subsequent analyses without reprocessing aligned reads a second time (aka incremental build). This feature is highly relevant for large studies with incremental releases such as ENCODE and GTEx and also for individual lab projects where datasets or samples are added as the project evolves.

In the second step of the pipeline, the MAJIQ quantifier is executed. As in the original MAJIQ framework, splicing quantification is performed in units of LSVs. Briefly, an LSV corresponds to a split in gene splicegraphs coming into or out of a reference exon. Each LSV edge, corresponding to a splice junction or intron retention, is quantified in terms of its relative inclusion (PSI, Ψ *∈* [0, 1]) or changes in its relative inclusion between two conditions (dPSI, ΔΨ *∈* [*−*1, 1]). Given the junction spanning reads observed in each LSV, MAJIQ’s Bayesian model results in a posterior distributions over the (unknown) inclusion level (ℙ (Ψ)), or the changes in inclusion levels between conditions (ℙ (ΔΨ)). This model accounts not only for the total number of reads but also for factors such as read distribution across genomic locations and read stacks. Given its Bayesian framework, the model can also output the confidence in inclusion change of at least C (ℙ (*|*ΔΨ*| > C*)), or the expectation over the computed posterior distributions (𝔼 [Ψ], 𝔼 [ΔΨ]). In this work, we introduce two new algorithms within the MAJIQ quantifier. The first involves how intron retention is quantified, allowing for much faster execution with higher accuracy (see Methods). The second addition is the implementation of additional test statistics, termed MAJIQ HET (heterogeneous). Conceptually, the original MAJIQ model assumes a shared (hidden) PSI value for a given group of samples and accumulates evidence (reads) across these samples to infer PSI. In contrast, MAJIQ HET quantifies PSI for each sample separately and then applies robust rank-based test statistics (TNOM, InfoScore, or Mann-Whitney U). As we demonstrate below, the new HET test statistics allow MAJIQ to increase reproducibility in small heterogeneous datasets, and gain power in large heterogeneous datasets.

A new optional analysis step introduced here is the VOILA Modulizer, an algorithm which organizes all identified LSVs into AS modules and then groups these modules by type. Briefly, AS modules represent distinct segments of a gene splicegraph involving overlapping LSVs which are contained between a single source and single target exon. However, unlike DiffSplice’s AS modules (*5*), we do not use a recursive definition of these modules and instead classify all identified modules by their substructures into types. The module’s substructures are in turn defined by the basic units of alternative splicing, namely intron retention, exon skipping and 3’ or 5’ splice variations. As we demonstrate below, the automatic AS module classification greatly facilitates a wide range of downstream analysis tasks.

The next step of the pipeline involves visualization of the quantified PSI and dPSI using VOILA v2. This new package runs as an app (on macOS, Windows, Linux) which supports the visualization of thousands of samples per LSV as violin beeswarm plots with multi group comparisons and advanced user filters. Users can perform searches by gene name or junction, and simplify the visualization by filtering out lowly included junctions. This option is highly relevant for large heterogeneous datasets where many junctions might be captured but may not be relevant for specific comparisons/samples. Notably, unlike the builder filters described above, the VOILA v2 filters do not affect the underlying splicegraphs but only help declutter the visualization to aid in subsequent analysis. VOILA v2 has the option to run as a server to share results with collaborators while all of the pipeline’s results can also be exported into other pipelines as tab-delimited files and for automated primer design for validation using MAJIQ-SPEL (*6*).

### Performance evaluation

In order to assess MAJIQ HET, our new method for detecting differential splicing, we performed a comprehensive comparison to an array of commonly used algorithms using both synthetic and real data. We considered only algorithms capable of analyzing large datasets, including the original MAJIQ algorithm (upgraded with the v2 code-base to enable efficient data processing), rMATS turbo, LeafCutter, SUPPA2, and Whippet. Figure 2A shows processing time and memory when performing a multi-group, multi-sample comparison, typical for such datasets. In this case, we perform all pairwise comparisons between 10 tissue groups, and the number of samples in each group grows from 1 (10 total samples) to 6 (60 total samples). All algorithms are able to process such large datasets using only 0.5-4 GB of memory, an amount readily available on modern laptops. However, large differences exist in terms of running time, with SUPPA2 (55 hours) and Whippet (50 hours) taking substantially longer to analyze the larger dataset (6 samples per group, 60 total samples) compared to approximately 6 hours by rMATS, LeafCutter and MAJIQ v2.

**Fig. 2:**
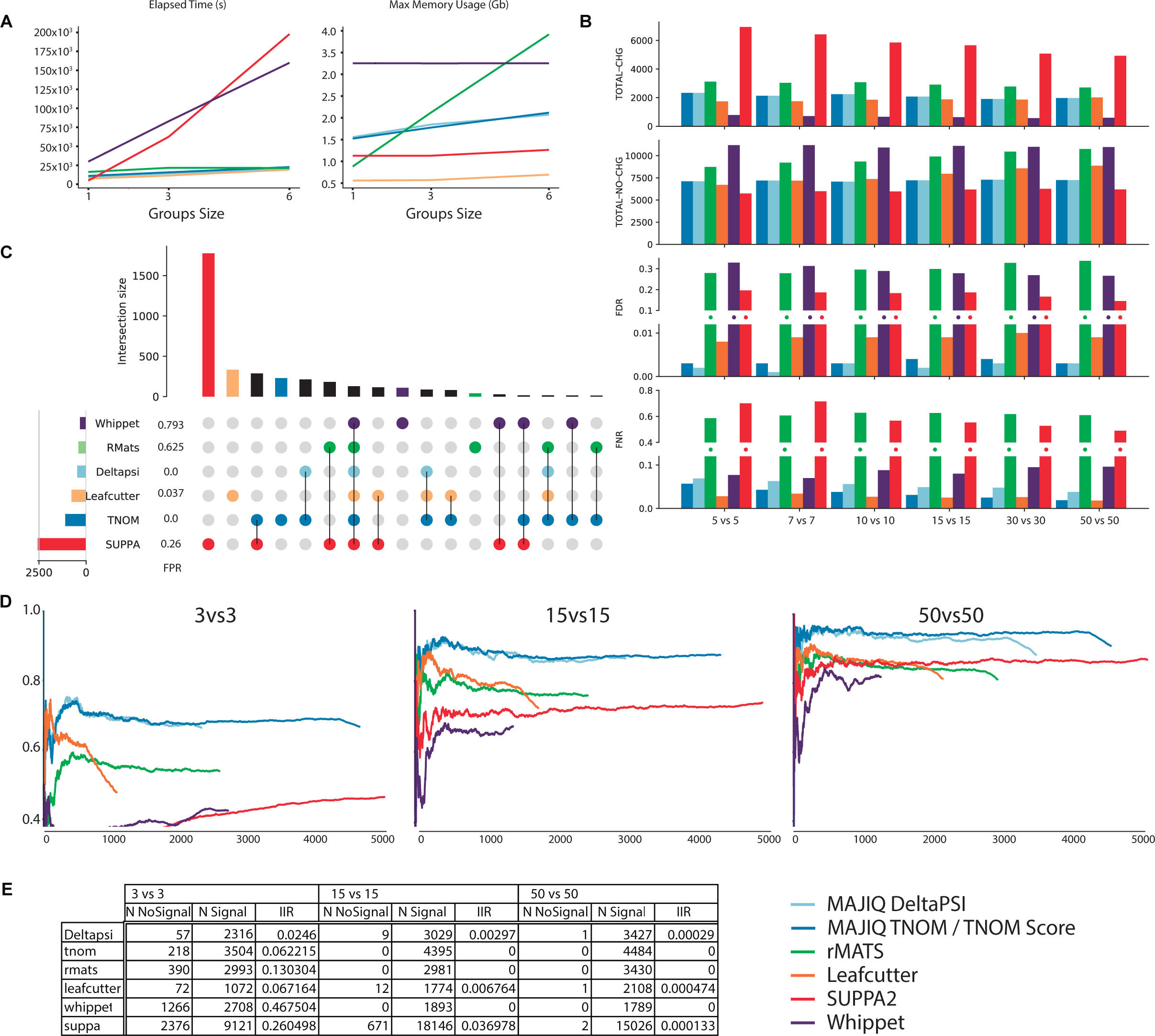
Performance evaluation using synthetic and real data. **(A)** Time (left) and memory (right) consumption when analyzing multiple sample groups. Results shown are for running all pairwise differential splicing analysis between 10 tissue groups from GTEx v8 as the number of samples per group increases from 1 to 6 (x-axis). **(B)** Performance evaluation for differential splicing calls using simulated GTEx cerebellum and skeletal muscle samples and aggregated over genes (see main text and Methods). Metrics include the total number of genes reported as changing or non changing by each method, and the associated FDR and FNR. X-axis denotes the size of the groups. **(C)** Upset plot based on the 10vs10 analysis shown in (B). The bars on top represent the overlap between genes reported as differentially spliced by each method indicated below it. The bars and FPR values by each method name on the left refer to genes reported only by that method. **(D)** Reproducibility ratio (RR) plots for real data, using GTEx cerebellum and liver samples. Analysis here is based on each method’s reported list of splicing events (not genes) and unique scoring approach. X-axis is the ranked number of events reported by each method and Y-axis is the fraction of those events reproduced within the same number of top-ranking events when repeating the analysis using a different set of samples from the same tissue groups. The length of the line represents the total number of differentially splicing events reported by each method (see Methods for details). RR graphs are shown for comparing group sizes of 3 (left), 15 (middle), and 50 (right). **(E)** Intra-to-Inter Ratio (IIR) results for GTEx samples as in (D). IIR computes the ratio between the number of events reported as significantly changing when comparing two sample groups of the same type (N No Signal column) and the number of events reported as significantly changing when comparing groups of different types (here GTEx liver and cerebellum samples as in (D)).

Next, we assessed the accuracy of all algorithms using a large-scale synthetic dataset for comparing two tissue groups. This synthetic dataset, by far the largest of its kind to the best of our knowledge, was constructed to be “realistic” such that each synthetic sample was generated to mimic a real GTEx sample from either cerebellum or smooth muscle tissues (see Methods). All methods were required to report changing AS events which pass the method’s statistical significance test and inferred to exhibit a substantial splicing change of at least 20% (see Methods). However, we note that since the various algorithms use significantly different definitions of AS events it is hard to compare those directly. For example, LeafCutter defines AS events as clusters of overlapping introns which may involve multiple 3’/5’ alternative splice sites and skipped exons, while rMATS is limited to only classical AS events with two alternative junctions. Thus, to facilitate a comparative analysis, we resorted to comparing the various algorithms output at the gene rather than event level using the synthetic dataset shown in Fig. 2B. A more refined analysis at the AS event level for each method can be found in Fig. S1 and follows the same trends discussed here at the gene level. First, we found SUPPA2 consistently reported over 6,000 differentially spliced genes, thousands more than any other method, while Whippet reported roughly 785 genes, significantly less than the other methods which reported over 2,000 changing genes (Fig. 2B top bar chart). Whippet, followed by rMATS, reported significantly more non-changing events. SUPPA2, rMATS, and Whippet all exhibited high FDR ranging around 15-30%, with the former two also exhibiting high FNR over 40%. Both MAJIQ and MAJIQ HET consistently maintained a lower false discovery rate compared to other algorithms (0.3%) and a low level of false negative rate which was similar to that of LeafCutter. On small sets, for example when using 5 samples per group, LeafCutter had a slightly lower FNR (2.5% vs 5.5% for HET), but MAJIQ exhibited lower FDR (0.03% vs 0.8%) while still reporting overall 34% more genes as changing (2,337 vs 1,739) and 6% more as non-changing (7,110 vs 6,713). It is also worth noting that the actual difference in the number of changing AS events reported by MAJIQ and LeafCutter is significantly higher, with 4,267 reported by MAJIQ vs. 2,169 by LeafCutter. This increased difference is mainly due to the increased resolution of event definition by MAJIQ. Specifically, MAJIQ uses the local splice variations formulation described above, while LeafCutter uses a definition of overlapping intronic regions which give rise to coarser event definition and can be sensitive to the coverage threshold used.

The significant differences between the methods described above raises the question how the reported sets of differentially spliced genes overlap. Fig. 2C illustrates the result of such analysis when using 10 samples per group. Here, we looked at the intersection between different methods at the gene level and when a set was unique to a method (i.e. the underlying events are well defined) we also estimated the associated FPR. We found SUPPA2 reports a significantly higher number of unique genes (1,777) as differentially spliced but over a quarter of those are false positives. The next set sizes are those for LeafCutter (333), HET and SUPPA2 (288), HET (230), and MAJIQ HET and PSI (214) with a FPR of 4% for the LeafCutter’s unique set and close to 0 FPR for both MAJIQ’s algorithms unique sets. rMATS and Whippet report significantly fewer unique genes with a high false positive rate of 62% and 79% respectively.

Next, we turned to assess performance on real GTEx data using several metrics. Here, unlike the synthetic data analysis which focused on comparative evaluation at the gene level, we focus on the actual AS events reported by each method. First, we used the reproducibility ratio (RR) statistic as shown in Figure 2D. The RR plots follow a similar procedure to that of irreproducible discovery rate (IDR) plots, used extensively to evaluate ChIP-seq peak callers (*3, 7*). Briefly, RR plots answer the following simple question: given an algorithm *A* and a dataset *D*, if we rank all the events that algorithm *A* identifies as differentially spliced (1*, …, N_A_*), how many would be reproduced if you repeat this with dataset *D^′^*, comprised of similar experiments using biological or technical replicates? The RR(*n*) plot, as shown in Fig. 2D, is the fraction of those events that are reproduced (y-axis) as a function of *n ≤ N_A_* (x-axis), with the overall reproducibility of differentially spliced events expressed as RR(*N_A_*) (far right point of each curve in Fig. 2D). In our RR analysis using groups of size 3 to 50 GTEx samples each, we found both MAJIQ and MAJIQ HET compared favorably to the other methods, but with the new HET algorithm exhibiting improved detection power resulting in a higher number of AS events at the same reproducibility level.

The second statistic we used for evaluating performance on real data is the intra-to-inter ratio (IIR) (*4*), which serves as a proxy for FDR on real data where the labels are unknown. Specifically, IIR computes the ratio between the number of differentially spliced events reported when comparing groups of the same condition (e.g. brain) and the number of events reported for similar group sizes of different conditions (e.g. brain vs liver). In our work, we found IIR to be a lower bound estimate of true FDR, though it lacks theoretical guarantees. In the analysis shown in Fig. 2E, we found IIR to behave similarly to FDR on synthetic data with MAJIQ, MAJIQ HET, and LeafCutter exhibiting low IIR of 2%-6% even for small group sets of 5 samples, while rMATS, SUPPA2, and Whippet had an IIR of 13%, 26% and 46% respectively. However, unlike FDR on synthetic data, IIR dropped much more significantly, hitting practically zero for all methods for large sample groups. This result is to be expected since the IIR statistic compares sample groups of the same type, unlike the synthetic dataset described above where different tissues are compared.

The last component we included for assessing different methods’ accuracy is a comparison to PSI quantifications using triplicates of RT-PCR assays, the gold standard in the RNA field. We previously produced over 100 such experiments from two different mouse tissues and showed MAJIQ compared favorably to SUPPA and rMATS (*3, 4*). Here, we extended this analysis to LeafCutter and found that MAJIQ’s quantifications correlates significantly better with those of RT-PCR (see Fig. S2). We note that this analysis for LeafCutter was possible since all events we tested were simple cassette exon skipping, but it is not clear how to translate LeafCutter’s output to actual PSI in the general case.

### VOILA v2 enables visualization of thousands of samples

To facilitate visualization and downstream analysis of both the new outputs from MAJIQ HET over large, heterogeneous datasets and traditional MAJIQ PSI or MAJIQ dPSI quantification over replicate experiments, we developed VOILA v2 as a server based cross-platform app. Replacing the previous HTML file based visualization with VOILA v2 allows for interactive visualization of all LSVs in all genes, with data ranging from one sample to thousands of samples. After an initial indexing step that is run one time, users can now, on the fly, filter their data by several criteria including dPSI levels between groups, read coverage over junctions, LSV types and complexity, and the statistical test for significance, as opposed to re-running VOILA with the filtering criteria, as was required in the previous version. Another advantage of the new VOILA v2 is the ability to run it as a server so that results can be shared with collaborators without the need to transfer large files.

To highlight these new features, we ran MAJIQ HET and VOILA v2 on GTEx v8 brain tissues which are known to exhibit high levels of alternative splicing. Overall, this analysis involved 2,335 RNA-seq samples from 374 donors across 13 tissue groups (see Methods). Figure 3A shows the VOILA view for this large dataset for the key splicing factor gene *PTBP1*, including a splicegraph (top) with combined read information from 225 cerebellum RNA-seq samples. Users can easily add and remove splicegraphs for other tissue groups or individual samples of interest. Figure 3A bottom panel shows a VOILA visualization for quantifying a single junction in a single LSV across the 2,335 RNA-seq samples. Here, the 13 tissues are displayed as violin beeswarm plots with each point representing a single sample which can be interrogated by hovering the user’s cursor over it. Finally, VOILA uses a heatmap (Fig. 3A bottom right) to represent the pairwise differences between the tissue groups for the junction of interest. The upper half of the heatmap represents the difference in medians of 𝔼 [Ψ] distributions between the tissue groups, while the bottom half represents the p-values associated with these group differences (see Methods). For the example LSV and junction in *PTBP1*, the cerebellar tissues (cerebellum and cerebellar hemisphere) show a distinct splicing pattern with reduced usage of this junction (lower 𝔼 [Ψ] values in the left-most violin plots) which was significant according to MAJIQ HET (InfoScore shown) (Figure 3A).

**Fig. 3:**
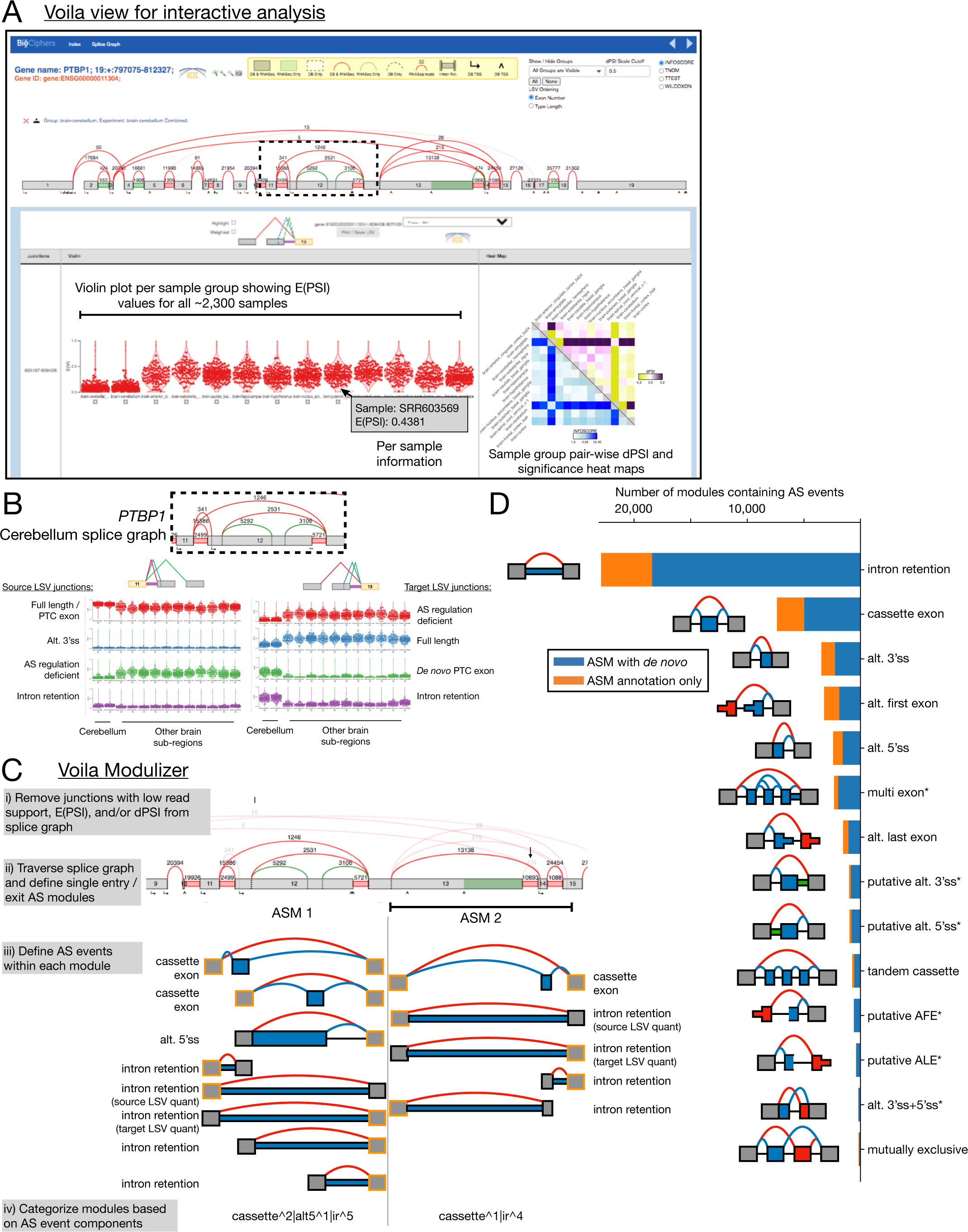
Enhanced visualization of large datasets and downstream analysis of alternative splicing modules with VOILA v2. **(A)** VOILA view of MAJIQ HET output for 13 brain tissue groups from GTEx from 2,335 RNA-seq samples originating from 374 unique donors. Top portion shows gene information and filtering criteria as well as the splicegraph for *PTBP1* showing combined reads from 225 cerebellum samples. Bottom portion displays visualization and PSI quantification for each junction in each LSV for the gene of interest. Here the distribution of 𝔼 [Ψ] values across the indicated tissue groups is displayed as a violin beeswarm plot for the red junction for the exon f13 target LSV, represented in the cartoon, for all 2,335 RNA-seq samples. Individual sample information is given by hovering the cursor over individual points that represent each sample (gray box). Bottom right heatmap displays MAJIQ HET quantifications of all group pairwise comparisons across the 13 brain tissue groups to highlight significant splicing changes. Yellow to purple color scale on the top right indicates the expected ΔΨ between tissue groups while blue color scale on the bottom left indicates the significance of the difference between group PSI distributions for one of four statistics used by MAJIQ HET (InfoScore displayed). **(B)** Top shows region of human *PTBP1* splicegraph (with reads from combined cerebellum samples) and two LSVs corresponding to a mammalian specific exon skipping event that alters PTBP1 splicing regulatory activity (*11*) (green junction in exon 11 source LSV, left; red junction in exon 13 target LSV, right) and *de novo* detection of a conserved, PTC-containing exon previously shown to be included in mouse neuronal tissues (*3*) (green junction in exon 13 target LSV). Bottom shows distribution of PSI across the 13 brain tissue groups as well as annotation of each junction. **(C)** VOILA Modulizer workflow (gray boxes) and an example region of the *PTBP1* splicegraph where junctions that did not meet a median 𝔼 [Ψ] value of 5% or more in any of the 13 brain tissue groups were removed (arrows). Two alternative splicing modules (ASMs) were defined as single entry, single exit regions of the splicegraph and within these modules binary, AS events are defined. Gray exons highlighted in yellow indicate reference exons that belonged to LSVs for which MAJIQ quantification exists. Blue junctions and exonic or intronic regions indicate inclusion of the alternative region of the event and red junctions indicate exclusion of the alternative region. **(D)** Stacked bar chart showing the number of binary AS event types that make up AS modules across the 13 brain tissue groups from GTEx. AS event types are represented with a cartoon to the right of the chart and are named to the left of bars. Asterisks indicate non-classical AS event types. Each junction or intron had to have a median of 𝔼 [Ψ] values of 5% or more across the samples of at least one tissue group to contribute to AS module definitions. Blue regions indicate AS events that contained *de novo* junctions and/or introns not found in the annotated transcripts (Ensembl v94) while orange regions indicate AS events containing only annotated junctions and introns.

### VOILA Modulizer defines alternative splicing modules to facilitate down-stream analysis

The LSV and junction showcased in the above example are of biological importance. PTBP1 is a widely expressed splicing factor that binds CU-rich sequences, but it is downregulated during neurogenesis which contributes to neuronal splicing patterns (*8–10*). Decreased activity of PTBP1 in neuronal tissues is attributed to numerous mechanisms, some of which involve splicing regulation of two cassette exons in the region highlighted in the *PTBP1* splicegraph (Figure 3A,B boxed regions) (*3, 11*), making differences between brain subregions of potential interest. Mammalian-specific, neuronal skipping of an alternative cassette exon in the linker region between the second and third RNA recognition motifs (RRMs) of *PTBP1* (exon 12 in the splicegraph) results in a protein isoform of PTBP1 with reduced repressive activity leading to altered splicing patterns during neuronal differentiation (*11*). Additionally, in mouse brain we previously described inclusion of a unannotated, premature termination codon (PTC) containing, cassette exon with conserved splice sites in humans that shows increased inclusion in mouse cerebellum (compared to brainstem and hypothalamus) and is developmentally regulated through murine cortex development (*3*). While LeafCutter analysis of *PTBP1* on all of GTEx failed to detect this event in human tissues, we find evidence of *de novo* splice junction reads corresponding to both the conserved 3’ and 5’ splice sites of this unannotated exon that we validated previously in mouse (Figure 3B), suggesting this exon is also included in human brain tissues.

This region of the splicegraph is complex, however, and is defined by overlapping LSVs each with multiple splice junctions and intron retention detected (Figure 3B: exon 11 source LSV, left; exon 13 target LSV, right). While the LSV formulation has several benefits, including accurate PSI quantification of complex splicing patterns involving more than two splice junctions (*3*), it is difficult for users to know which junction quantifications and combinations of junctions from different LSVs should be combined to define common alternative splicing (AS) events, like the cassette exons described above in *PTBP1*. Moreover, while certain annotated and *de novo* junctions may have sufficient read coverage for detection and quantification by MAJIQ, they can be very lowly included in a user’s condition(s) of interest. For example, several hundred reads across GTEx brain samples support the existence of the annotated, intron distal alternative 3’ss of exon 12 of *PTBP1*, but source LSV quantification of the relative usage of this junction is low across all samples (Figure 3B, left. Blue junction median PSI across samples of *<* 5% in all tissue groups). Such junctions add additional complexity to the splicegraph and may hinder definition of common AS event types across the transcriptome.

To overcome these limitations and to facilitate downstream, transcriptome wide analysis of common AS event types we developed the VOILA Modulizer (Figure 3C). First, users have the option to simplify the splicegraph to remove junctions that do not meet a threshold for raw read coverage, low inclusion levels across the input samples (𝔼 [Ψ]), and/or low relative splicing changes between input comparisons between sample groups (𝔼 [ΔΨ]) (Figure 3Ci). This helps remove junctions that do not meet a user’s desired threshold for biological significance and facilitate downstream event definitions, like the alternative 3’ss of exon 12 of *PTBP1* discussed above with low inclusion levels across all sample groups (blue junction in Figure 3B, left). Next the simplified splicegraph is traversed to define single entry, single exit regions of the splicegraph that we call alternative splicing modules (AS modules or ASMs), as shown for part of *PTBP1* (Figure 3Cii). Within each AS module, pattern matching is performed between the remaining exon and junction structure of the simplified splicegraph to each of 14 basic AS event types (Figure S3A). This process is illustrated in Figure 3Ciii for two AS modules within *PTBP1*. We note that this step can lead to some redundant event information (e.g. intron retention events sharing the same junction and intron coordinates, as in Figure 3C). Because these events are quantified from both sides through a source and a target LSV, the quantification in terms of PSI or dPSI between conditions may not agree and thus both are provided. Nonetheless, downstream filtering can ensure agreement when counting event types and defining changing events.

Running the VOILA Modulizer produces a number of files based on event types with a uniform structure containing coordinate and quantification for each sample group to facilitate downstream analysis on AS modules and AS event types of interest (Figure S3A). Some AS event definitions identified by the Modulizer are analogous to those defined by other splicing quantification algorithms that only handle binary, classical splicing events (e.g. MISO (*12*) or rMATS (*13*)). However, the MAJIQ + VOILA Modulizer approach adds a number of benefits compared to other available algorithms. First, our approach allows for *de novo* splice junction and intron retention detection, which is crucial in the context of GTEx brain subregions. Using a simplification threshold of median 𝔼 [Ψ] over brain tissue groups of *≥* 5% to be included in the simplified splicegraph, we defined 32,435 AS modules where 70.6% contain at least one unannotated splice junction and/or intron retention (Figure 3D, Figure S3B). The AS module formulation also allows for definition of common splicing patterns across brain subregions beyond binary splicing events, which made up 59.2% of all AS modules. The remaining 40% of AS modules contained multiple AS events which, in many cases, involved mixing of a classical event type with intron retention (Figure S3B). Both at the AS event level (Figure 3D) and at the AS module level (Figure S3B), intron retention was particularly common using our simplification threshold of median 𝔼 [Ψ] of greater than 5% in any one brain tissue group. This is consistent with previous studies that have found neuronal tissues to have very high levels of intron retention compared to other contexts (*14*).

Initial analysis of the most common AS module types led us to add additional splicing event patterns to our definitions, beyond those that are classically defined in other tools (intron retention, cassette exon, alternative 3’ and 5’ss, alternative first and last exons, tandem cassette exons, and mutually exclusive exons (*12, 13*)). These included putative alternative first and last exons, where at least one alternative exon is created from a *de novo* junction that does not belong to any nearby exon, and putative alternative 3’ or 5’ss, where a cassette exon has an inclusion junction removed during simplification (low inclusion) with sufficiently high intron retention levels (see Figure S3A for full details). These new splicing event types participated in the make up of 13.1% of AS modules or 11.6% of AS events overall detected in the brain (Figure 3D, event types marked with asterisks). Importantly, the Modulizer outputs all of these event types in a format amenable to downstream regulatory analysis, which will facilitate the future characterization of these splicing patterns (Figure S3A).

### Analysis of unique cerebellar splicing patterns highlights known and novel regulatory programs

Finally we wished to use MAJIQ + VOILA Modulizer to analyze differential splicing patterns between brain subregions. Previous studies focused on splicing quantitative trait loci within GTEx brain tissues found the cerebellar tissues cluster separately from other brain subregions based on splicing (*15*). Our analysis of *PTBP1* (Figure 3B) and pairwise analysis of the number of significant LSVs according to MAJIQ HET further supports distinct splicing patterns in cerebellar tissues (Figure S4A). For these reasons we sought to identify AS modules and events with unique splicing patterns in the cerebellum. Using the above AS module definitions from all junctions and introns with group level median 𝔼 [Ψ] *>* 5%, we next searched for consistent splicing changes between the two cerebellar tissues (cerebellum and cerebellar hemisphere) and other brain subregions using MAJIQ HET. We required an absolute difference in median 𝔼 [Ψ] values of 20% or more when comparing both cerebellar tissue groups to the same other brain region tissue group in addition to having a Wilcoxon rank-sum *p <* 0.05 (Figure 4A, see Methods).

**Fig. 4:**
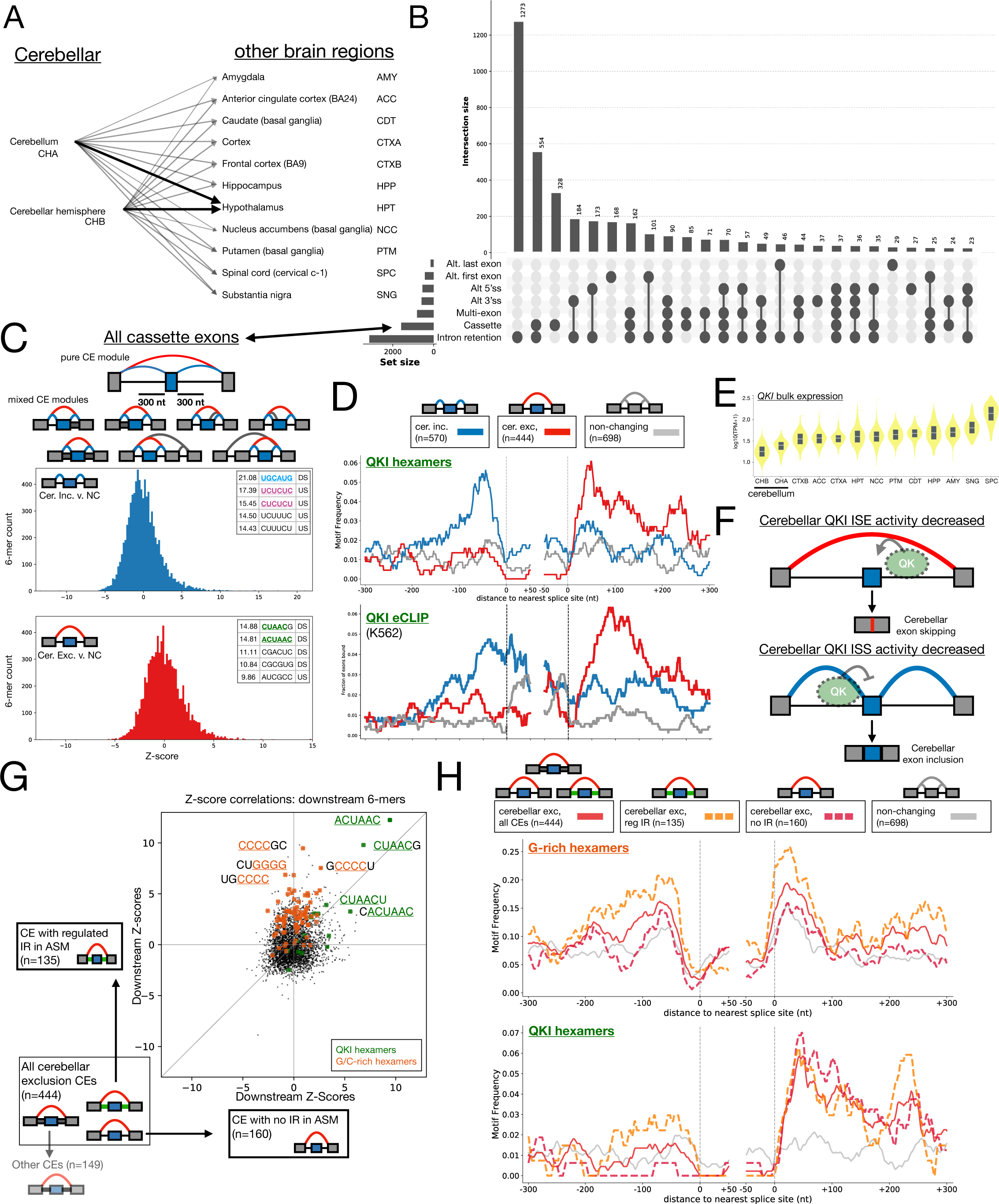
MAJIQ HET + VOILA Modulizer defines the complex landscape of cerebellar splicing changes and regulation. **(A)** Pairwise comparisons run through MAJIQ HET to find significant splicing changes between GTEx cerebellar tissues (cerebellum and cerebellar hemisphere) versus the other 11 brain tissue groups. Dark arrows indicate an example of a consistent change, where both cerebellar tissue groups versus the same other brain region, the hypothalamus, shared a significant change. Alternative splicing modules (AS modules) were kept for downstream analysis if at least one such consistent comparison was significant (see Methods). GTEx abbreviations are given to the left of each tissue. **(B)** Upset plot showing the consistent, significantly changing AS event type(s) that make up AS modules. AS events had to have an absolute difference in median 𝔼 [Ψ] of 20% or more when comparing both cerebellar tissue groups (cerebellum and cerebellar hemisphere) to the same other brain region tissue group in addition to having a Wilcoxon rank-sum *p <* 0.05 as reported by MAJIQ HET. **(C)** Top shows examples of all cassette exons (CEs) used for motif analysis. All CEs have quantified inclusion junctions (blue junctions) and a shared exclusion junction (red junction), potentially within a mixture of other AS event types (gray junctions and introns). Bottom shows distribution of Z-scores for hexamer motif occurrences within 300 nucleotides upstream or within 300 nucleotides downstream of all CE events when comparing CEs showing significant cerebellar inclusion versus CEs that did not change when compared to other brain regions (see Methods) (middle, blue) or when comparing CEs showing cerebellar exclusion versus non-changing CEs (bottom, red). Top motifs corresponding to putative binding sites of RBPs of interest are highlighted (QKI (green), RBFOX (light blue), SRRS6 (yellow), SRSF11 or PTB (purple)). All motifs and Z-scores are given in Table S1. **(D)** RNAmaps showing the frequency of QKI hexamer motif occurrence (top, ACUAAY frequency over sliding windows of 20 nucleotides with smoothing using a running mean of 5 nucleotides) or in vivo binding of QKI (K562 eCLIP peaks frequency, bottom) around cerebellar inclusion (blue), exclusion (red), or non-changing (gray) CEs. **(E)** *QKI* bulk tissue gene expression (log_10_ (1 + TPM) for ENSG00000112531.16) sorted by median brain tissue expression. Chart generated using `gtexportal.org`. **(F)** Model for QKI position dependent regulation in GTEx brain tissues. Decreased expression of QKI in cerebellar tissues results in a decrease in downstream intronic splicing enhancer (ISE) activity of QKI, leading to cerebellar exon exclusion (top, red), and a decrease of upstream intronic splicing silencer (ISS) activity of QKI, leading to cerebellar exon inclusion (bottom, blue), when compared to other brain tissue groups. **(G)** Scatter plot showing hexamer Z-score correspondence for two non-overlapping sets of cerebellar CE exclusion events: (y-axis) CE exclusion events which came from AS modules containing changing intron retention (IR) event(s) versus non-changing and (x-axis) CE exclusion events from AS modules without IR event(s) detected. Motifs of interest are highlighted according to colors in the inset. **(H)** RNAmaps of motifs of interest for given sets of cerebellar exclusion cassette exon event sets. Top shows G-rich hexamer motif occurrence (five of six positions are G and contains GGGG) while bottom shows QKI hexamers (ACUAAY). Frequencies were determined over sliding windows of 20 nucleotides with smoothing using a running mean of five. Lines indicate CE exon set according to the legend: red, all cerebellar exclusion CEs; orange dashed, subset of exclusion CEs which also contained a changing IR event; fuchsia dashed, subset of exclusion CEs with no IR event with the AS module; gray, all CEs which were not changing between comparisons.

From these comparisons we found 3,995 unique, changing AS modules (Figure 4B) comprising over 7,500 changing AS events (Figure S4B). At the changing AS module and AS event levels, intron retention was prevalent, followed by cassette exons and other mixtures of binary AS event types with intron retention (Figure 4B). As with the analysis based on inclusion levels alone (Figure 3), most changing AS modules (53.3%) consisted of multiple, binary AS event types (Figure 4B), highlighting the prevalence of complex splicing changes and the power of MAJIQ + VOILA Modulizer approach.

Alternative splicing regulation of cassette exons in neuronal tissues is very well studied with a number of expression changes associated with splicing factors (e.g. expression of the RBFOX family, down regulation of PTB proteins, expression of NOVA proteins, etc.) (*9, 10*). For this reason we wished to analyze the regulatory signature around the cassette exons defined from our MAJIQ HET + VOILA Modulizer analysis to see if we could capture known, and potentially novel, regulatory motifs around cerebellar cassette exons.

Our initial analysis focused on all changing cassette exon (CE) events. This mirrors the CE landscape that would be identified by other splicing quantification algorithms and consists of a combination of CEs which come from modules consisting of only a single CE event in addition to those from complex modules with multiple event types (Figure 4B, arrowhead, Figure 4C, top). Because RNA binding proteins bind short motifs and splicing factor binding that results in alternative splicing regulation typically occurs proximal to the splice sites of an alternative exon (*16*), we performed a Z-score analysis for hexamer occurrence within 300 nucleotides upstream or downstream of cerebellar changing cassette exons versus those alternative exons that did not change between brain subregions (see Methods). Moreover, because splicing factors typically act in position-specific manners (e.g. binding downstream of a cassette exon enhances exon inclusion while binding downstream represses inclusion) (*17, 18*), we further separated cassette events into those with increased exon inclusion in cerebellar tissues (Figure 4C, blue) and those with increased exon exclusion in cerebellar tissues (Figure 4C, red) when compared to other brain subregions.

Supporting the validity of our approach, this analysis uncovered a number of motifs either upstream or downstream of cerebellar cassette exons with known links to neuronal splicing regulation. For example, for cerebellar inclusion cassettes we found a number of CU-rich and UGC containing hexamers upstream and the RBFOX-binding-motif, UGCAUG (*19*), enriched downstream (Figure 4C, blue). SRRS6/nSR100 is known to bind UGC-containing sequences upstream of neuronal microexons to enhance their inclusion with the aid of SRSF11 that binds CU-repeat sequences (*20*). Accordingly, motif maps across our different cerebellar exon classes based on hexamers shown to bind SRRS6 (*21*) and SRSF11 (*20*) by iCLIP show clear enrichment of these motifs just upstream of cerebellar inclusion cassette exons. This result is consistent with increased expression of these two genes in cerebellar tissues leading to enhanced intronic splicing enhancer (ISE) activity around these events (Figure S5A-E).

In addition to SRRS6 and SRSF11, the RBFOX family is highly expressed in neuronal tissues and is known to enhance exon inclusion when it binds downstream of the 5’ss (*22, 23*) (Figure S5G). Indeed, we find a strong enrichment of the known UGCAUG-binding site just downstream of cerebellar inclusion events (Figure 4C, blue). This result is consistent with increased expression of these genes and increased ISE activity in the cerebellum versus other brain subregions (Figure S5F-H).

Interestingly, we found hexamers containing motifs known to bind QKI (e.g. ACUAA containing (*24*)) were enriched around both cerebellar inclusion (upstream) and exclusion events (downstream) (Figure 4C). QKI is known to act as a splicing enhancer when it binds downstream of cassette exons and represses exonic inclusion when it binds upstream (*24*). We generated a motif map of the QKI hexamer (ACUAAY (*25*)) around these exon classes and found clear positional enrichment proximal to the regulated splice sites in both exon sets (Figure 4D, top). Moreover, we generated RNA maps of in vivo binding events (determined by CLIP peaks) of QKI across multiple cell types and found enriched binding consistent with the motif maps (Figure 4D, bottom, Figure S5I). Compared to other brain subregions, the two cerebellar tissues exhibited lowest expression of QKI (Figure 4E). This result points to a regulatory mechanism by which decreased expression of QKI in cerebellum may contribute to both cerebellar exon exclusion events (loss of enhancing activity downstream leading to exon skipping) and cerebellar exon inclusion events (loss of repressive activity upstream leading to inclusion) (Figure 4F).

Given that many regulated cassette exons occur within AS modules containing other AS event types (Figure 4B), we next wished to explore if regulatory motifs differed between these subsets. Because AS modules containing cassette exons and those containing both cassette exon and intron retention events are common (Figure 4B, Figure S3B, S4B), we chose to stratify the set of all cassette exons into those that contained a regulated intron retention event and those that occurred in AS modules in which intron retention was not detected. We calculated Z-scores for hexamers from these exon subsets by comparing them against the set of exons that were not changing in cerebellar comparisons and compared the results of the two analyses. Figure 4G shows an example of this analysis for hexamers located downstream of cerebellar exclusion cassette exon subsets. The top two hexamers that match QKI binding motifs (ACUAAC and CUAACG) found when analyzing all CE events (Figure 4C) also had the highest Z-scores in the intron retention regulated and no intron retention CE subsets (Figure 4G, green). On the other hand, several of the G- and C-rich motifs that were enriched downstream of all CE cerebellar exclusion events (Figure 4C) were biased towards higher Z-scores solely in the CE subset that contained regulated intron retention (Figure 4G, orange). This is consistent with observations from previous studies analyzing intron retention events that found retained introns tended to be more G/C-rich when compared to non-retained introns (*14*). Motif maps across the different cerebellar exclusion CE sets supported the Z-score analysis and highlight that the enrichment of G-rich sequences (Figure 4H, top) and C-rich sequences (Figure S6A,B) around all cerebellar exclusion CEs is driven mostly by the subset of CEs containing a regulated intron retention event (compare dashed orange and dashed fuchsia lines). The QKI hexamer showed similar positional enrichment downstream of both CE subsets (Figure 4H, bottom).

Similar results were seen when comparing Z-scores for upstream and downstream hexamers identified in the all cassette exon analysis (Figure 4C) of cerebellar inclusion and exclusion CE subsets stratified by intron status (Figure S6A). While some of the motifs found in the complete CE analysis scored similarly in subsets stratified by intron retention status (e.g. the RBFOX hexamer or SRRS6 hexamers around cerebellar inclusion exons), others showed biased enrichment in CEs with regulated intron retention compared those with no intron retention (e.g. CU-repeat hexamers) (Figure S6). Overall, this analysis highlights some shared and distinct regulatory features of cerebellar cassette exons with and without evidence of intron retention.

## Discussion

The work presented here represents the culmination of continuous development of MAJIQ since its original release in 2016 (*3*). The original MAJIQ, like many other algorithms, was designed for comparing relatively small groups of RNA-seq from biological replicates. However, as we demonstrate here using GTEx v8, datasets nowadays can easily grow to hundreds and thousands of non-replicate samples. The sheer size and heterogeneous nature of such data poses challenges that go beyond just algorithm efficiency. Additional challenges include the ability to capture but also simplify *de novo* and complex splicing variations, the ability to define subtypes over such complex splicing events, and the ability to visualize and process such events and subtypes for downstream analysis. MAJIQ v2 is the only algorithm, to the best of our knowledge, that supports such features through efficient implementation of several algorithmic innovations we introduced here: The simplifier, the modulizer, incremental build options, and the VOILA v2 visualization package. In addition, we perform extensive comparison of MAJIQ v2 to other algorithms, create a resource for reproducible algorithm comparison in the form of both data and software package, and demonstrate the utility of the new splicing analysis features by performing a detailed analysis of differential splicing between more than 2,300 samples from GTEx v8 brain subregions.

The algorithmic contributions in this work include a new method to quantify *de novo* intron retention, an incremental build, addition of the MAJIQ HET statistics which do not assume a shared PSI between samples in a group, and the modulizer in VOILA. The resulting new features enhance splicing analysis, especially on larger datasets. For example, MAJIQ’s incremental build saves much of the processing needed when adding new samples to existing repositories. Labs or centers can thus process data such as GTEx once, then efficiently add more relevant samples as needed. Notably, our performance evaluations discussed below show performance for the first analysis, but subsequent analyses can be expected to be even faster. Furthermore, as these datasets get larger, we also expect to see more *de novo* junctions. These junctions increase the complexity of the splicegraph and the size of splicing events considered. The MAJIQ simplifier enables users to more finely control how this complexity enters the analysis.

The new features of MAJIQ v2 are accompanied by matching ones in VOILA v2 visualization and analysis package. The VOILA Modulizer provides a new view of splicing changes on the splicegraph in terms of splicing modules. These complex units can be broken down into classical splicing events that may share similar splicing regulation, but allow for a more refined classification than traditional approaches, as demonstrated in our analysis of brain subregions. In contrast, tools that only list classical events (e.g. rMATS) quantify those solely based on the reads within these event definitions. Consequently, reads outside these event definitions, which can greatly alter the splicing quantification, are ignored. In addition, the VOILA viewer now allows for interactive visual analysis and supports a server version, allowing large analyses performed on cluster or cloud environments to be viewed without downloading large datasets locally.

In terms of performance we showed MAJIQ v2 compares favorably to available methods. In terms of efficiency, we showed MAJIQ v2 is as fast and memory efficient as the two most efficient tools, rMATS-turbo and LeafCutter. This is a notable achievement given that MAJIQ is the only tool amongst those that offers detection and quantification of *de novo* intron retention. Accounting for IR in splicing analysis is computationally expensive but nonetheless important in many settings as we discuss below.

On synthetic data, MAJIQ and LeafCutter were the only two tools that simultaneously demonstrated both low FDR and FNR when identifying genes with differential splicing. We note that our usage of LeafCutter included additional filtering for ΔΨ *>* 20% beyond the default p-value based filtering as we found that the default settings performed much worse (*26*). Whippet was the only other tool that also exhibited low FNR, but it demonstrated FDRs over 20%. Our results suggest that many genes called as differentially spliced by Whippet, rMATS, and SUPPA are false discoveries. Furthermore, it suggests that rMATS and SUPPA miss a substantial fraction (*>* 40%) of the genes that it should call as differentially spliced.

On real RNA-seq data from GTEx we found MAJIQ outperformed the other tools. Specifically, MAJIQ’s reproducibility, measured using the reproducibility ratio, was consistently higher than all other tools. The difference between MAJIQ and other tools was particularly striking when comparing a small number of samples but persisted even when comparing 50 vs 50 samples. Comparing MAJIQ HET introduced here to MAJIQ dPSI from (*3*), we found both to have similar reproducibility, but HET offered a significant increase in detection power. While LeafCutter was comparable to MAJIQ on the synthetic dataset, we found that its reproducibility on real data was not, exhibiting reproducibility lower than rMATS and comparable to SUPPA2. When using intra-to-inter ratio (IIR) to assess false discovery, we found IIR approached 0 when considering larger numbers of samples for all tools. However, for very small sample numbers of 3 vs 3, only MAJIQ and LeafCutter achieved IIR below 10%.

The extensive evaluations we performed here serve not just to assess the specific tools we included, but as a service for the community. First, we created the largest synthetic RNA-seq dataset to date, with over 300 samples. In contrast to many other works, the data generated here was based on real life GTEx samples. It also does not reflect MAJIQ’s model and was based instead on transcript-based quantifications by other algorithms (RSEM). As such, we would expect it to benefit tools that are built around a similar model (e.g. SUPPA). A second contribution is the evaluation package we created, validations-tools. This package allows users to not only reproduce our results but also to easily add future tools and repeat the analysis for future developers or for anyone who wants to assess performance on their own unique dataset. We highly recommend researchers and cores to take advantage of this as it is possible that on a dataset with other characteristics the various algorithms would perform differently. Finally, we note that the efforts to create reproducible results in genomics and specifically for tool development are constantly ongoing. We previously documented in detail issues we identified with using using outdated software, software misuse, and lack of reproducibility for analysis scripts and data that severely affected software assessment, including MAJIQ (*26*). We hope the reproducibility tools we included here will help avoid such issues and make it easier for future developers to achieve at least the “bronze” level of reproducibility as was recently proposed (*27*).

Finally, applying our improved pipelines to GTEx brain subregions allowed us to define the complex alternative splicing patterns observed across over 2,300 heterogeneous human neuronal tissue samples from 374 donors and 13 tissue groups. Our approach and subsequent analysis offers several advances compared to previous efforts. For example (*15*) also analyzed differential splicing in brain subregion splicing but included only annotated, classical splicing events identified by rMATS. Several other GTEx analyses use LeafCutter’s framework and focus on detecting sQTLs. Our work advances these efforts through improved quantification accuracy (described above) and our LSV based approach, which is the only method able to capture *de novo* and complex splicing events as well as retained introns (IR). Furthermore, as we illustrated here for cerebellum specific regulation, our newly introduced definition of AS modules and AS event types greatly facilitate downstream regulatory analysis.

Applying MAJIQ HET and AS subtypes from the VOILA modulizer allowed us to discover additional, novel complexity within transcripts for the crucial splicing regulator, PTBP1, including a *de novo*, premature stop codon containing exon in human that we previously validated in mouse brain subregions (*3*). This exon was preferentially included in cerebellar tissues, leading us to focus on the cerebellar specific splicing program. Our regulatory analysis on cerebellum specific cassette exons highlighted many known splicing regulators previously shown to be essential in neuronal splicing programs (i.e. the RBFOX family, SRRS6 with SRSF11, PTBP1, and QKI (*9, 10, 20*)), highlighting the validity of our approach and definitions of cassette exons based on complex LSVs. Crucially, the MAJIQ + VOILA Modulizer approach allowed us to stratify this superset of cassette exon events into different subsets based on the presence or absence of other AS event types within the module (e.g. CEs with or without intron retention). While some motifs are shared and similarly enriched around CEs with and without regulated intron retention (e.g. RBFOX and QKI), other motifs were specifically enriched in the intron containing subset only. In the case of cerebellar exon exclusion events, the signal for G/C rich motifs observed on the superset of all CEs was driven entirely by the subset of CEs containing intron retention events. We anticipate the new ability we introduced here to interrogate AS modules made up of combinations of AS event types will facilitate future regulatory discoveries in other datasets from additional biological contexts.

We note that there are key limitations to the regulatory analysis we performed for cerebellar-specific splicing, which was based solely on bulk tissue RNA-seq experiments from GTEx. Previous work leveraging single cell data to deconvolute bulk GTEx tissues into their relative cell type compositions suggests that cerebellar tissues contain relatively larger proportions of neurons compared to other brain subregions (*28*). This fact can confound the interpretation of our results in terms of neurobiology as neurons are known to express certain splicing factors (e.g. RBFOX3/NeuN, SRRS6), which may explain the cerebellar splicing pattern we observed here. Thus, future directions for improving MAJIQ involve accounting for cell type heterogeneity as well as combining long reads for isoform specific deconvolution. Other promising directions for future exploration include analysis of RNA sequencing for clinical diagnostics and exploiting MAJIQ’s advantages for improved sQTL analysis.

In summary, we introduced here a significant update to the original MAJIQ package. MAJIQ v2 empowers fast, detailed, and accurate analyses of large heterogeneous RNA-seq datasets and is already supporting a highly active user group spanning hundreds of labs, centers and companies across the world. Our analysis of brain subregions provides a compelling example of such analysis on over 2,300 human neuronal tissue samples leading to several novel findings related to cerebellum specific splicing regulation. We hope the analysis we performed, along with the tool, data, and evaluation package we supply, will inspire many more researchers to delve into splicing regulatory analysis in their own data and make exciting new discoveries.

## Methods and Materials

### MAJIQ builder

In this subsection, we review how the MAJIQ builder prepares the structure and observations per experiment that are used for downstream splicing quantification as part of a scalable and principled approach to splicing analysis of large numbers of experiments. We describe the MAJIQ builder’s new approach for estimating intron read rates, which allows junction and intron coverage to be calculated once and reused efficiently for multiple analyses, unlike other methods that quantify intron retention. We also describe the MAJIQ simplifier, which reduces the complexity of the structural models of splicing used in quantification that especially arises from the analysis of large and heterogeneous datasets.

MAJIQ encodes the set of all possible splicing changes for a gene in terms of a splicegraph. A splicegraph is a graph-theoretic representation of a gene’s splicing decisions from one exon to another, with exons as vertices and junctions and retained introns as distinct edges connecting exons. The exons of each gene are non-overlapping genomic intervals. Each junction has a source and target exon with a position within each exon, indicating the positions that are spliced together when the junction is used. Retained introns are between adjacent exons and indicate that intron retention between the exons is possible.

MAJIQ first constructs each gene’s splicegraph by parsing transcript annotations from a GFF3 file. Exon boundaries and junctions from each transcript for a gene are combined in order to produce the minimal splicegraph that includes each transcript’s annotated exons and junctions, splitting exons by retained introns to ensure that each junction starts and ends in different exons. MAJIQ then updates the splicegraph with *de novo* junctions and introns found from processing input RNA-seq experiments’ junction and intron coverage.

MAJIQ processes aligned input RNA-seq experiments to per-position junction and intron coverage in the following way. First, MAJIQ identifies reads with split alignments. The genomic coordinates of each split corresponds to a potential junction. Meanwhile, the coordinate of the split on the aligned read is the junction’s “position” on the read. MAJIQ counts the number of reads for each junction from each possible position. Afterwards, MAJIQ identifies reads that contiguously intersect known or potential introns (i.e. reads that intersect the genomic coordinates between adjacent exons without splits within the intron boundaries). If the intron start is contained in the aligned read, the intron “position” is defined as for junctions (treating the exon/intron boundary as a junction with zero length). For aligned reads intersecting the intron but not the start, additional positions are defined by the genomic distances of the first positions of the aligned reads to the intron start. These additional positions per intron increase the number of ways aligned reads can intersect introns in comparison to junctions. To adjust for this and model intron read coverage similarly to junction read counts, MAJIQ aggregates together adjacent intron positions to the equivalent number of possible positions per junction, taking the mean number of reads per reduced positions.

MAJIQ uses the obtained junction and intron coverage to update the splicegraph in the following way. Each potential junction is mapped to matching genes by prioritizing (1) genes that already contain the junction (i.e. annotated junctions) over (2) genes where both junction coordinates are within 400bp of an exon, which are prioritized over (3) genes where the junction is contained within the gene boundaries. The input experiments are divided into user-defined build groups. MAJIQ adds a *de novo* junction to the splicegraph if there is sufficient evidence for its inclusion in one of the build groups. This happens when the total number of reads and total number of positions with at least one read exceeds the user-defined minimum number of reads and positions in at least a minimum number of experiments. MAJIQ adds new *de novo* exons or adjusts existing exon boundaries to accommodate the added *de novo* junctions as previously described. Potential introns are added to the splicegraph under similar criteria, and their boundaries are adjusted or split to accommodate the adjusted or *de novo* exon boundaries. Since processed intron coverage is averaged over the entire original intronic region, we can carry over the same coverage as an estimate for all resulting splicegraph introns, which are contained in the original intron’s boundaries. In contrast, MAJIQ’s previous approach, which is also used by most other tools that quantify intron retention, quantified intron coverage using local counts of unsplit reads sharing the position of known junctions. These local counts must be calculated using information from all processed experiments (for all *de novo* junctions), which requires samples to be reprocessed each time an analysis with different samples are performed. MAJIQ’s new approach allows intron coverage to be processed once and used for multiple builds with potentially different intron boundaries. This enables MAJIQ’s new incremental build feature, which saves intermediate files with junction and intron coverage that can be calculated once and reused instead of BAM files for multiple builds. This reduces storage and time processing experiments that are part of multiple analyses.

While MAJIQ uses raw totals of read rates and number of nonzero positions for adding junctions and introns to the splicegraph, the MAJIQ builder performs additional modeling of per-position read rates for use in quantification. First, we mask positions with zero coverage and with outlier coverage. Outlier coverage is assessed under the observation that per-position read rates generally follow a Poisson distribution. For each junction/position, we use all other positions with nonzero coverage for that junction to estimate the Poisson rate parameter. Then, MAJIQ calls any position with an extreme right-tailed p-value (default 10*^−^*^7^) under this model an outlier and ignores its contribution to coverage for quantification. Second, we perform bootstrap sampling of the total read rate over unmasked positions in order to model measurement error of true read rates. Under the assumption that each unmasked position is identically distributed, MAJIQ performs nonparametric sampling with replacement to draw from a distribution with identical mean and variance as the observed positions (see supplementary note). Since we assume that our read rates are generally overdispersed relative to the Poisson distribution, MAJIQ replaces nonparametric sampling with Poisson sampling when the nonparametric estimate of variance is less than the mean (i.e. underdispersed).

MAJIQ performs quantification of splicing events modeled as LSVs, which are defined by a splicegraph. A source (target) LSV is defined for an exon as a choice over the incoming (outgoing) edges to (from) that exon from (to) a different exon. In general, only LSVs with at least two edges are considered. MAJIQ builder prepares output files with raw and bootstrapped coverage for each junction/intron in each LSV for quick use by downstream quantifiers.

We observed that builds from many build groups or with high coverage tend to have increasingly complex splicegraphs and LSVs with many junctions. Many of these junctions are often lowly used in all the samples but were included in the splicegraph because they had enough raw reads and positions (noisy *de novo*) or are part of an unused annotated transcript. This motivated the MAJIQ simplifier, which allows junctions and introns to be masked from the final splicegraph used for quantification. After the splicegraph is constructed using all input build groups, MAJIQ calculates the ratio of the raw read rate for each junction/intron relative to the other junctions/introns in each LSV. If a junction has consistently low coverage in each of the build groups relative to the other choices in the two LSVs it can belong to, it is “simplified” and removed from the final splicegraph. This reduces the complexity of the final splicegraph and quantified LSVs, making output files smaller and downstream quantification more efficient.

In summary, the MAJIQ builder combines transcript annotations and input RNA-seq experiments in order to build a splicegraph encoding all possible splicing events consistent with both annotations and data and to prepare read coverage for quantification in terms of LSVs. The MAJIQ builder’s new approach for estimating intron read rates allows junction and intron coverage to be calculated once and reused as part of an incremental build for multiple analyses, unlike other methods that quantify intron retention. The MAJIQ builder also introduces an approach for simplifying the complexity that arises in splicing events when processing large numbers of experiments. Overall, this allows the MAJIQ builder to produce structural models of possible splicing events and read coverage for downstream quantification that scale to the setting of large numbers of RNA-seq experiments.

### MAJIQ quantifiers

MAJIQ provides three methods for quantifying RNA-seq experiments. MAJIQ PSI, MAJIQ dPSI, and MAJIQ HET, which we introduce in this paper. MAJIQ PSI and dPSI, which were previously described in (*3*), quantify groups of experiments that are assumed to be replicates with a shared true value of PSI per group. MAJIQ PSI estimates a posterior distribution of PSI (Ψ) for a single group, while MAJIQ dPSI compares these distributions for two groups in order to estimate a posterior distribution for dPSI (ΔΨ). MAJIQ HET compares two groups of samples but drops the replicate experiments assumption, enabling analysis of more heterogeneous samples. Instead, experiments are quantified individually and groups are compared under the assumption that the true values of PSI are identically distributed between the two groups.

All three pipelines share the same underlying machinery for inferring posterior distributions for Ψ. Formally, Ψ for a junction in an LSV is defined as the fraction of expressed isoforms using the junction out of all expressed isoforms containing the LSV. This fraction is not directly observable. Instead, we observe the number of reads aligned *r_j_* to each junction *j* in the LSV. We model each *r_j_* as a realization of a binomial distribution over the isoforms with probability Ψ*_j_* :

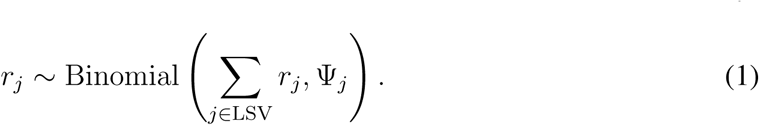

We take a Bayesian approach to integrate prior knowledge of Ψ, allowing for improved estimation when there is low read coverage. This requires a prior distribution on Ψ. We previously observed that most values of Ψ are nearly zero or one, which can be modeled using a generalization of the Jeffrey’s prior for an LSV with *J* junctions:

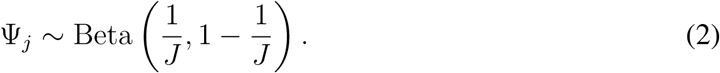

This prior is conjugate to the binomial likelihood, allowing for efficient closed-form estimation of the posterior distribution of Ψ*_j_* given the observed number of reads:

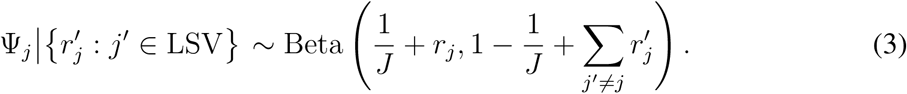

Since MAJIQ build obtains bootstrap replicates of observed read rates, we perform this posterior inference on each set of bootstrap replicate read rates to obtain an ensemble of posterior distributions.

For MAJIQ PSI, we obtain this ensemble of posteriors for replicate experiments by adding the observed read rates from the experiments that pass more stringent reads and position thresholds than the builder. MAJIQ PSI treats the average of the posterior distributions as a final distribution over Ψ. It reports point estimates of Ψ as the mean of this distribution (𝔼 [Ψ]) and saves a discretized version of the distribution for visualization in VOILA.

MAJIQ dPSI takes this a step further by using the posterior distributions on Ψ_1_, Ψ_2_ for two groups in order to compute ΔΨ = Ψ_2_ *−* Ψ_1_ between the two groups. We start by computing the distribution of ΔΨ under the assumption of independence of Ψ_1_ and Ψ_2_ by marginalizing the product of their distributions:

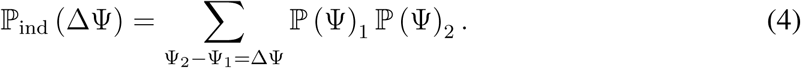

not independent, so we integrate our knowledge that ΔΨ is usually close to zero as a prior on ΔΨ. Following our previous work, we formulate our prior ℙ_prior_ (ΔΨ) as a mixture of three components: (1) a spike around ΔΨ = 0, (2) a broader centered distribution around ΔΨ = 0, and (3) a uniform slab. We determine our final posterior distribution on ΔΨ by adjusting ℙ_ind_ (ΔΨ) by the prior and renormalizing:

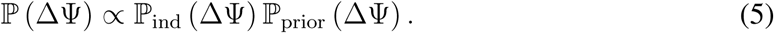

MAJIQ dPSI computes point estimates of ΔΨ using the posterior mean of the distribution (𝔼 [ΔΨ]) and identifies confidence of measured changes in inclusion as posterior probabilities ℙ (*|*ΔΨ*| > C*).

MAJIQ HET takes a different approach for comparing inclusion between two groups of experiments. MAJIQ HET drops the assumption of replicate experiments to consider heterogeneity in Ψ between experiments within a group. Instead, MAJIQ HET assumes that the values of Ψ per experiment in each of the groups come from the same distribution. We evaluate this assumption using null hypothesis significance testing. Null hypothesis significance testing is performed using one (or more) of four tests: (1) Welch’s two-sample t-test, (2) Mann-Whitney U test, (3) Total Number of Mistakes (TNOM) test, and (4) InfoScore test. Welch’s two-sample t test and Mann-Whitney U test are well-documented elsewhere (*29, 30*). Our implementation of Mann-Whitney U test computes exact p-values when there are at most 64 experiments and computes asymptotic p-values using normal approximation with tie and continuity correction for larger samples. Meanwhile, the InfoScore and TNOM tests are adapted from ScoreGenes (*31*). The TNOM test evaluates how well a single threshold on PSI can discriminate between the observed values in the two groups. The Total Number of Mistakes is the minimum number of misclassified observations under the best possible thresholds. The distribution on TNOM when the distributions are equal are calculated using the closed-form formula in (*32*) to obtain p-values. Similarly, the InfoScore test evaluates how well a single threshold discriminates between groups, but, instead of measuring misclassifications directly, it identifies the threshold with the highest mutual information between the threshold and the true group labels. MAJIQ HET uses the dynamic programming algorithm in (*32*) to evaluate the distribution of InfoScore under the null hypothesis in order to obtain p-values. All four tests require observed values of Ψ per experiment, which is not directly observed. MAJIQ HET accounts for variable uncertainty per experiment in our estimations of Ψ by repeated sampling of Ψ from the posterior distributions of quantified samples. MAJIQ HET computes the p-value for each repeated sample of Ψ over quantified experiments and reports the 95th-percentile over the resulting p-values. These p-value quantiles are not calibrated, so MAJIQ HET also computes p-values with the posterior means of Ψ. MAJIQ HET also reports the median of the observed posterior means of Ψ for each group. These p-values and the difference between the median observed posterior means are used together downstream in VOILA for the identification of high-confidence differentially spliced LSVs.

### VOILA

VOILA provides a suite of post-processing and visualization tools designed to allow researchers to make use of MAJIQ quantifications directly, or easily format and filter the output for passing to other post-processing tools.

The VOILA viewer acts as a complete visualization tool for interactive analysis of output from MAJIQ PSI, dPSI, or HET. It includes search and filter mode for all discovered LSVs, as well as an in-depth viewer for the full splicegraph of a gene and all of the LSVs found within it. When using the VOILA viewer with output from MAJIQ HET, VOILA will also automatically generate heatmaps for each LSV with the to quickly indicate the discovered ΔΨ and statistical results from each group comparison. The viewer frontend runs completely within a web browser interface, so it is able to function with similar results on any modern operating system without installation of special frameworks or system libraries. The viewer can also be configured to run as a standalone web server such that the interactive results can be easily shared with collaborators. Tutorials and parameters are made available to integrate VOILA with a wide range of common web server production software.

VOILA also has a number of modes for filtering and rearranging data into a number of human and machine-readable files. Determining confidently non-changing (background) and confidently changing events is one of the primary use cases. We define highly-confident non-changing events from MAJIQ HET as being (1) above a nominal p-value threshold, (2) within-group variance is sufficiently low as measured by IQR, and (3) between-group ΔΨ is sufficiently low as measured by difference in medians. We accept that the between-group ΔΨ threshold may be redundant in combination with the other two thresholds. We define confident changing events from MAJIQ HET as being (1) below a p-value threshold and (2) between-group ΔΨ is sufficiently high as measured by difference in medians.

In addition to the basic text output modes, there is a separate comprehensive output mode dedicated to finding specific event types/patterns called the VOILA Modulizer. The VOILA Modulizer searches for a large number of relevant patterns, both common and complex.

Each set of events is delimited on the basis of AS “modules” found by MAJIQ in each analyzed gene. Modules refer to areas of the splicegraph between single entry (one junction path, diverges to two or more) and single exit (all junction paths converge back to one).

Inside each of the AS “modules” detected by the modulizer, smaller AS “events” (sub patterns matching specific known organizations of junctions or introns) are then categorized. Currently, the list of potential patterns we match to find an event is fixed to a specific set, which can be found in Figure S3. All events which do not match any known splicing pattern are dumped to an “other” category which may be of possible interest in rare cases.

Modulizer supports any number or combination of MAJIQ experiments as input, in the form of PSI, dPSI, and/or HET VOILA files. These are used for narrowing modules to form around junctions / introns we find relevant, as well as to verify which AS modules and AS events are changing or non-changing, based on coverage, Ψ, and differences in Ψ (ΔΨ). All filters may be disabled or adjusted.

At a high level, Modulizer uses a sequential pipeline for filtering and assembling output. First, all junctions and introns are read, and any which do not pass the reads, PSI, and/or dPSI thresholds are immediately removed from consideration. Then, using the remaining introns, and junctions, Modulizer identifies AS modules by looking for genomic locations with single-entry / single-exit as previously described. Then, Modulizer filters and removes modules which do not pass criteria such as not being sufficiently changing, lack of LSVs, or being constitutive. After filtering, Modulizer performs pattern matching for each AS event type on each AS module to identify all component AS events. Finally, Modulizer scans the input VOILA files for relevant quantifications in order to produce output TSV files for each individual AS event type, a high-level summary of all events found in each discovered module, and a summary of quantifications per module suitable for generating a heatmap according to the user’s filtering criteria (e.g. the shortest discovered junction within the AS module to represent the inclusive AS product, the most changing junction in the AS module from HET and/or dPSI inputs, etc.).

### Sample selection from GTEx

We selected from GTEx in the following way. We required all samples to have a RIN score of greater than 6. For performance evaluation we chose to evaluate a comparison between cerebellum and skeletal muscle. We randomly selected 150 samples from both tissues, excluding the same donor from being selected in both tissues. For the brain subregions analysis, we selected all samples in GTEx v8 associated with brain tissue (not including pituitary gland). We also performed another analysis with all tissues in GTEx v8 using 30 or less samples per tissue. Samples were downloaded as FASTQ or as BAM and converted to FASTQ depending on when they were released. Samples that were part of v7 are available on SRA, so they were downloaded using SRA Tools (v2.9.6) as FASTQ files. New samples from the v8 release were only available as BAMs on the cloud, so they were downloaded and converted to FASTQ using samtools (v1.9).

### Simulated RNA-seq as ground truth

We used the expression quantification data from the GTEx v8 release as the basis for our simulations. Briefly, we downloaded publicly available gene- and transcript-level quantification tables for GTEx v8 from the GTEx portal (https://www.gtexportal.org/home/datasets). To match how the GTEx consortium performed these analyses, we downloaded the GRCh38 build of the reference genome sequence and gene models from v26 of the GENCODE annotation.

We selected 300 samples from GTEx to serve as the basis for 300 simulated samples, each real sample providing the expression distribution underlying one simulated sample. To run BEERS, we first need to prepare four configuration files that are customized for the desired dataset: geneinfo, geneseq, intronseq, and feature quants. The geneinfo, genenseq, and intronseq files define the structure and sequence information for each simulated transcript. As a result, these three files are determined solely by the choice of reference genome build and annotation. The feature quant files are specific to each individual sample and define a distribution of transcript-level expression. First, we used the genome sequence and gene models to create the geneinfo, geneseq, and intronseq files. Since the genome is fixed across all simulated samples. We used the same set of these files to simulate all GTEx-derived samples. Next, we extracted TPM values for each sample from the GTEx transcript quantification table and used these distributions of TPM values to generate separate BEERS feature quant config files for each simulated sample. Lastly, to determine the total number of reads to simulate for each sample, we used the gene-level quantification file to count the total number of gene-mapping reads in each GTEx sample.

To simulated strand-specific reads with uniform coverage across no errors, substitutions, or intron retention events, we ran the BEERS simulator using the following command-line options: `-strandspecific -outputfq -error 0 -subfreq 0 -indelfreq 0 -intronfreq 0 -palt 0 -fraglength 100,250,500`.

We transformed ground-truth transcript abundances into ground-truth splicing quantifications for each splicing quantification tool, taking into account the tools’ differing definitions of splicing events. First, we defined ground-truth abundances for each exon or junction by adding the abundances of all transcripts including the exon or junction. Then, for each tool, we adopted their splicing event definitions, mapping the exon/junction abundances to compute their splicing quantifications.

### MAJIQ

MAJIQ reports splicing quantifications with respect to LSVs. Therefore, ground-truth values for PSI were calculated by dividing the ground-truth abundance of each junction by the sum of the ground-truth abundances for all junctions in each LSV.

### rMATS

rMATS reports a different format file per event type. But since all of them are classical binary event types, all can be reduced to two paths events, inclusion and exclusion. Each file contains the exon that defines each of the ways, so we calculate the Ψ_gt_ as inclusion/(inclusion + exclusion) using the exon transcript combination to get the exons ground-truth abundances for all junctions in each LSV.

### LeafCutter

LeafCutter reports splicing quantifications with respect to intron clusters composed of several junctions. Ground-truth values for LeafCutter’s splicing ratios were calculated using ground-truth junction abundances, similar to MAJIQ.

### SUPPA2

SUPPA2 reports classical events similarly to rMATS. So the approach we use here is similar to that tool. The main difference is that SUPPA2 reports the junctions coordinate in each one of the paths, so we use those junctions ground truth quantification to obtain the Ψ_gt_ as inclusion / (inclusion + exclusion).

### Whippet

Whippet outputs a psi.gz that contains the psi quantification of an event. That PSI is their formulation of the quantification from inclusion and exclusion paths. Differently to SUPPA2 or rMATS, Whippet combines a set of junctions to define a path, emulating in that way a transcript (or a portion of it). So, in order to find Ψ_gt_ of those paths, we look for those transcripts that include all the junctions (and virtual junctions). We combine the expression of those transcripts to find the Ψ_gt_ of each path.

### RNA-seq sample preprocessing before splicing analysis

We aligned RNA-seq reads from real and simulated GTEx samples to the human genome for splicing analysis with MAJIQ and other tools using the following procedure. Simulated GTEx samples were generated as pairs of FASTQ files. We performed quality and adapter trimming on each sample using TrimGalore (v0.4.5). Some tools require reads aligned to the genome. For these tools, we used STAR (v2.5.3a) to perform a two-step gapped alignment of the trimmed reads to the GRCh38 primary assembly with annotations from Ensembl release 94. Other tools required transcript quantifications relative to annotated transcripts. For these tools, we used Salmon (v0.14.0) using the trimmed samples to estimate transcript abundances.

### Performance evaluations

We wrote a package of evaluation scripts, called validations-tools, in order to compare MAJIQ in terms of speed, memory footprint, accuracy, and reproducibility for each one of the following tools: rMATS, LeafCutter, SUPPA2, and Whippet. This package was written to allow future users to not only reproduce our results but to easily add future tools and repeat these kinds of analyses with different datasets.

We adjusted the tools parameters following recommendations by each tool’s authors. Specific parameters are listed in Table S2. For these comparisons, we evaluated the methods’ computational efficiency and ability to identify splicing differences.

First, we evaluated computational efficiency of the different methods. We evaluated computational efficiency in terms of runtime and peak memory usage. Not all tools provide an extensive log of their execution, so, in order to measure wall time and memory usage, we used the output of `‘/usr/bin/time -v‘`. We ran each method for all pairs comparisons between 10 groups with increasing sample sizes on an Ubuntu Linux environment with 32 cores (Intel Xeon 2.7GHz and 64GB RAM).

Second, we evaluated the different methods’ performance in quantifying splicing differences on simulated and real datasets. On the simulated datasets, where we know ground-truth differences in splicing between transcripts, we calculated true and false positive rates for the identification of splicing differences by each method. However, on real datasets, where no ground-truth is available, it is not possible to calculate true or false positive rates. Instead, we evaluated two metrics, reproducibility ratio (RR) and intra-to-inter ratio (IIR), on real (and simulated for comparison) data. The first metric, RR, measures the internal consistency of differential splicing tools. This internal consistency is reflected in the assumption that each tool should identify roughly the same events when repeating a comparison between two groups using different samples. We quantify this by performing two such comparisons and computing the fraction of the top *n* differentially-spliced events in the first comparison that are also in the top *n* events of the second comparison. This produces a “reproducibility-ratio” curve, RR(*n*) for the method as a function of the number of top events. If the first comparison yields *N* “significant” events, RR(N) is called the reproducibility ratio. For the specific case of MAJIQ, we note that in order to comparisons of LSV-type events more comparable to classic AS events such as used by rMATS, we filtered out overlapping LSVs (i.e. those that share junctions) in order to avoid double-counting classic AS events. For example, a classic exon-skipping event would have matching source and target LSVs that overlap. However, we note that this filtering only reduces *N_A_* but does not affect the reproducibility curves (apart from extending to a different value of *N_A_*) (Fig. S7). Although reproducibility of a method on real data is a scientifically important goal, it is not a sufficient goal because highly biased methods can be highly reproducible. To address this limitation, the second metric, IIR, is based on the principle that comparisons between (inter-) two groups should have many more significant events than comparisons within (intra-) a group. Furthermore, significant events within the group are likely false positives. This is quantified by computing the ratio of the number of significant events from an intra-group comparison to the number of significant events from an inter-group comparison. We evaluated these metrics for each tool with varying sample sizes to identify which methods outperformed each other in different settings.

### Event-level evaluations

In these evaluations we check reproducibility and accuracy of reported differentially spliced events by the various tools shown in Figure 2. As we describe in the main text, each tool defines alternative splicing events differently so that direct comparison of the events or their number between tools is not possible. Thus, when using real data each method was assessed by its own set of reported events to compute reproducibility ratios (RR) and intra-to-inter ratio (IIR) as in Figure 2D,E.

In contrast, when using GTEx based simulated data we do have the “ground truth” (denoted “gt” below) for the abundance of each transcript. We thus use these values to summarize Ψ and ΔΨ observed in each method reported AS events and assess accuracy using the following definitions:

- True Positive: max ΔΨ_tool_ *≥* 20% and pvalue_tool_ *≤* 0.05 and max ΔΨ_gt_ *≥* 20%
- True Negative: max ΔΨ_tool_ *<* 5% and pvalue_tool_ *>* 0.05 and max ΔΨ_gt_ *<* 5%
- False Positive: max ΔΨ_tool_ *≥* 20% and pvalue_tool_ *≤* 0.05 and max ΔΨ_gt_ *<* 5%
- False Negative: max ΔΨ_tool_ *<* 5% and pvalue_tool_ *>* 0.05 and max ΔΨ_gt_ *≥* 20%
- Ambiguous: all other cases (when either ΔΨ *∈* [5%, 20%) or when ΔΨ and pvalue reported by the tool conflict), where max is taken over all junctions/introns that belong to each AS event.

The above definitions were used to assess accuracy at the event level for each method, as shown in Figure S1, and also served as the base for gene level evaluations described below.

### Gene-level evaluations

To facilitate more direct comparison between the different methods shown in Figure 2 we aggregated each tool AS events and their respective annotation as TP, TN, FP, and FN as given above to assess gene level performance. Naturally, gene level labels of TP, TN, FP and FN are defined based on the events they contain. The gene level labels are easy to define as positive or negative when all AS events embedded in it are considered positive or negative respectively. The problem arises when a gene has some of its events as false positives and false negatives. In that case, we prioritize the labels according to the following order: FP, FN, TP, TN. This means for example that an occurrence of a false positive event in a gene (according to the method’s specific event definition) would be counted as a false positive gene even if some other events were correctly labeled as true negative or even true positives. The rationale for this prioritization is that (a) positive events are expected to be rare and (b) we care the most about trying to validate or follow up on wrong hits (false positives) followed by missing true changes (false negatives).

### GTEx brain subregion analysis

#### MAJIQ HET and VOILA Modulizer on brain subregions

MAJIQ HET was run on all 78 unique pairwise comparisons of GTEx v8’s 13 brain tissue groups, and the results were visualized with VOILA. Significant LSVs were those considered to be those containing at least one junction or intron with an absolute difference in group median 𝔼 [Ψ] values of 20% or more between the two tissue groups and all four HET statistics (Wilcoxon, InfoScore, TNOM, and t-test) with *p <* 0.05.

The VOILA Modulizer was run on the resulting outputs with the following options: `--decomplexify-psi-threshold 0.05` to remove all junctions and introns from the splicegraph that had tissue group median E(PSI) of less than 5% across samples for every group; `--show-all` to include all AS modules and AS events in the output, not just those meeting the changing criteria. Default values were used for other options that flag changing AS modules and AS events in the output. For changing: a minimum absolute median difference in PSI between groups of 20% or more for the primary threshold and a p-value of less than 0.05 across all four MAJIQ HET statistics (Wilcoxon, InfoScore, TNOM, and t-test). For non-changing: a maximum absolute median difference in PSI of 5% or less between groups; a maximum interquartile range in PSI within a group of 10% or less; and a p-value of 0.05 or greater across the MAJIQ HET statistics).

#### PSI based AS module and AS counts across the brain

Counting of AS modules based on the initial PSI simplification across the 13 brain tissue groups was done by parsing the resultant VOILA Modulizer summary file. This file is organized by AS module and lists the number of each of the 14 AS event types, outlined in Figure S3A, contained in each. AS modules were classified and counted based on the presence or absence of each of the 14 AS event types. Certain AS event type definitions overlap. Specifically, every tandem cassette exon containing AS module will also contain a multi exon skipping AS event and every putative 5’ or putative 3’ss AS module will also contain an intron retention event. In these cases, the additional, partially redundant AS event type was added to the AS module classification if and only if their count within the module was larger than the count of the AS event they overlap with. For example, for an AS module to be classified as containing both tandem cassette exon (TCE) and multi exon skipping events (MES), the number of MES events within the module must be greater than the number of TCE events.

### Cerebellar AS module and AS event definitions

Given the large number of LSV-based splicing differences between the two GTEx cerebellar tissues (cerebellum and cerebellar hemisphere) and the other brain subregions according to MAJIQ HET comparisons (Figure S4A), we wished to define AS modules and AS events based on these comparisons. These two cerebellar tissues were derived from sampling in duplicate, with cerebellar hemisphere sampled during initial tissue collection (frozen) and cerebellum sampled after the brain was received at the brain bank (PAXgene) (*33*). Therefore, we focused our analysis on AS modules and AS events that displayed changes between both cerebellar tissues and one of the other subregions. For example, a cassette exon AS event would have to be labeled as changing according to the VOILA Modulizer filters (minimum absolute median difference in PSI between groups of 20% or more for the primary threshold and a p-value of less than 0.05) in both cerebellum versus cortex and cerebellar hemisphere versus cortex to be counted. We defined all such consistent, changing cerebellar AS events from the 14 AS event files output by the VOILA Modulizer and used these to count the number of modules containing each AS event type or combination of types.

### Cerebellar cassette exon regulatory analysis

To perform regulatory analysis around exons with differential cerebellar inclusion patterns we first defined a high confidence set of cassette exons (CEs) by applying additional filters to those described above. In addition to the primary filter of an absolute median difference in PSI of 20% or more between a cerebellar tissue and another brain subregion for one junction in the CE event, a secondary threshold of an absolute median difference in PSI of 10% or more was enforced for all four junction quantifications of the CE (i.e. the inclusion source LSV junction quantification, the inclusion target LSV junction quantification, and the shared exclusion junction quantified in both the source and target LSV). Next we enforced that the direction of change between the two exclusion junction quantifications and the two inclusion junction quantifications agreed in their direction of change in cerebellar versus other tissues. If both inclusion junction qualifications increased in cerebellar tissues and both exclusion junction quantification decreased, this was considered a cerebellar inclusion CE events. The opposite directions were considered cerebellar exclusion CE events. Non-changing CE events were defined as those flagged as non-changing by the VOILA Modulizer in every comparison of both cerebellar tissues versus the other 12 brain tissues. For CE subset analysis, CE with intron retention (IR) events were those where one or more of the CE junctions was also involved in a changing IR event in cerebellar versus other tissues. CE with no IR events were those CE events that came from modules without any IR events detected.

For sequence analysis we extracted GRCh38 sequences for intronic regions 300 nucleotides (nts) upstream and 300 nts downstream of every CE in each set. We calculated Z-scores by comparing the occurrence of each hexamer in the upstream intronic region in each cerebellar set of regulated CEs versus the non-changing set of CEs. This was repeated for the downstream intronic region as well.

Motif maps were generated to visualize position specific enrichment of particular hexamers of interest. Each hexamer, or set of hexamers, were searched for over sliding windows of 20 nts in the splice site proximal regions around the CE (i.e. intronic region 300 nt upstream of the 3’ss plus 50 nt downstream and 50 nt upstream of the 5’ss plus the intronic region 300 nt downstream). The frequency of occurrence was determined in each CE set and plotted using a running mean of 5 nts for smoothing.

RNAmaps for CLIP based binding of QKI were plotted in a similar way over the same splice site proximal regions. BED narrowPeaks files were downloaded for ENCODE eCLIP data (*34*) from encodeportal.org for QKI in K562 cells (accession ENCSR366YOG) or QKI in HepG2 cells (accession ENCSR570WLM) and replicate files were concatenated. BED narrowPeaks for uvCLAP data for QKI-5 in HEK293 cells (*35*) were downloaded from GEO (accession GSE85155) and lifted over from GRCh37 to GRCh38. These peak coordinates were overlapped with CE splice site proximal regions and the frequency of occurrence was assessed over the various cerebellar CE event sets at each position proximal to CE splice sites.

## Acknowledgments

We would like to thank Dr. Elizabeth J. Bhoj for her input on the final manuscript and co-advising JKA during the project.

## Funding

National Institutes of Health grants R01 AG046544, R01 LM013437, R01 GM128096 (YB). National Institutes of Health grant F30 HD098803 (JKA). Blavatnik Family Fellowship in Biomedical Research (MRG).

## Author contributions

YB conceived the project. JVG, JKA, SSN, and YB developed and tested the methodology for MAJIQ HET. JKA conceived and implemented the final methodology for quantification, sampling, and statistical testing in MAJIQ HET. JVG did the initial work porting Python code from MAJIQ v1 to MAJIQ v2. JVG and YB developed the new IR quantification algorithm. JKA formalized the approach for bootstrapping readrates. JVG and JKA implemented the updated MAJIQ Builder and MAJIQ PSI. JVG implemented MAJIQ dPSI. CJG and PJ implemented the VOILA viewer with input from MRG, CMR, and AJ. PJ, CMR, AJ, and YB developed the methodology for VOILA Modulizer. PJ implemented the VOILA Modulizer. NFL and GRG generated the simulated RNA-seq data for the performance comparisons. JVG carried out the performance comparisons vs other tools. MRG and CR performed the modules analysis. MRG conceived and carried out the brain subregions analysis. YB, JKA, and MRG wrote the final manuscript with input from PJ and JVG. All authors read and approved the final manuscript.

## Competing interests

The MAJIQ software used in this study is available for licensing for free for academics, for a fee for commercial usage. Some of the licensing revenue goes to Y.B and members of the Barash lab.

## Data and materials availability

The code for MAJIQ and VOILA are available for academic/non-commercial use at majiq. biociphers.org. Licensing information for commercial use can be found at majiq. biociphers.org/commercial.php. The code for validations-tools will be made available at bitbucket.org/biociphers/validations-tools. All processed data and code to reproduce figures will be deposited in a Zenodo before publication.

GTEx data used for the analyses in this manuscript are available in dbGaP under accession phs000424.

## Supplementary Materials

### Supplementary Note: procedure for bootstrapped readrates from perposition reads

MAJIQ’s bootstrapping procedure can be defined as follows. Without loss of generality, consider a single junction. For each RNA-seq read aligned with a split for this junction, we define the read’s position relative to the junction (or vice-versa) with a position *i* and count the number of reads associated with each position, which we call *S_i_*.

These raw readrates include biases that we would like to correct for; in particular, we define an explicit procedure for removing stacks by comparing the number of reads at each position against a Poisson model using the observed readrates at all other positions, which results in a set of stack-corrected nonzero readrates *R_i_* for *i ∈ {*1*, …, P}*, where *P* is the number of nonzero positions after stack removal. These are the observed units for bootstrapping, so to emphasize:

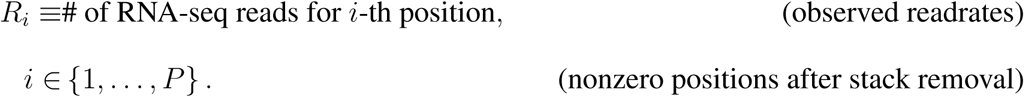

Other methods typically sum directly over positions *R_i_* (really *S_i_* since they generally also ignore read stacks) to produce a total junction readrate for use in quantification:

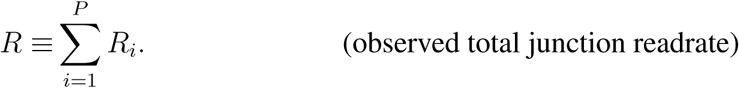

Since we are unsatisfied with uncertainty/variance accounted for by directly using *R*, we generate samples from a bootstrap distribution over the *P* nonzero positions.

If we make the assumption that we are given the number of nonzero positions *P* and that the underlying readrate for each of these positions is independent and identically distributed with finite mean 𝔼 [*R_i_*] = *µ* and variance 𝕍 [*R_i_*] = *σ*^2^, we can derive the mean and variance of our observed total readrate:

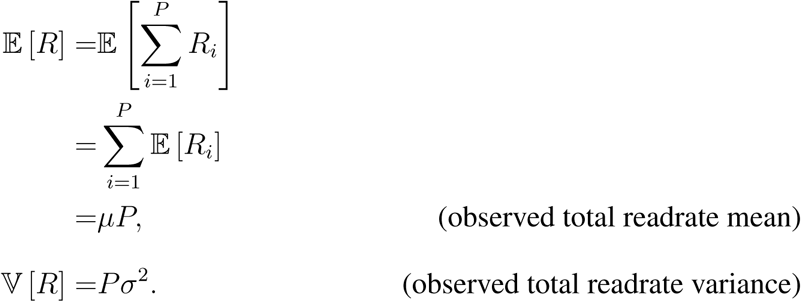

If we were able to take two samples for the observed total readrate (i.e. *R* and *R^′^*), their difference has mean 0 and variance 2*Pσ*^2^.

We define our bootstrapping procedure over observed nonzero reads *R*_1_*, …, R_ℙ_* to generate bootstrapped total reads *R̂*, *R̂’*, … such that the variance of the difference between bootstrap samples would be equivalent to that of the difference between two samples from the true distribution (i.e. 2*Pσ*^2^). In order to do this, we take *P −* 1 samples from *{R*_1_*, …, R_P_ }* with replacement and scale their sum by *P/*(*P −* 1).

It is straightforward to see that the bootstrapped total readrate has the same mean as the observed total readrate. In order to prove that the variance of the difference between two sample matches, we note that the covariance Cov *R_Z_k__, R_Z_k_′_* between any two draws from the observed per-position readrates with *Z_i_ ∼* Uniform(*P*) is:

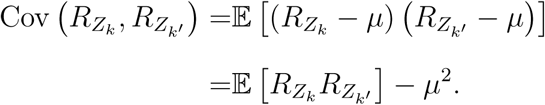

. We note that 𝔼 [*R_i_R_j_*] = *σ*^2^*δ_ij_* + *µ*^2^ (where *δ_ij_* is the Kroencker delta). When *k* = *k^′^*, it follows that 𝔼[*R_z_k__*,*R_z_k’__*] = *σ*^2^ + *μ*^2^. Otherwise, the law of total expectation gives:

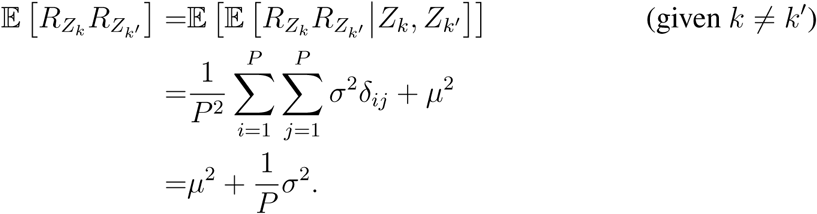

Combining the two cases, we have:

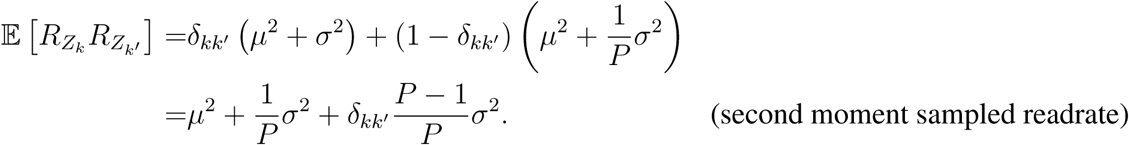

Therefore,

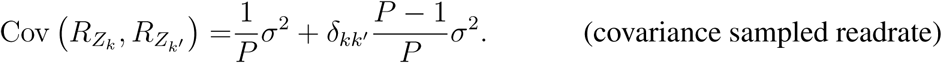

Thus, the variance of the bootstrapped total readrate is

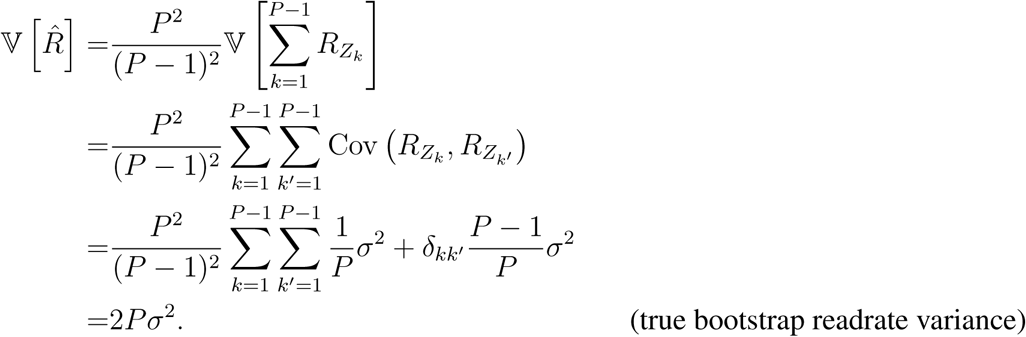

But we want the variance of the difference between two samples from the bootstrap procedure.

So, we calculate the covariance between two distinct samples:

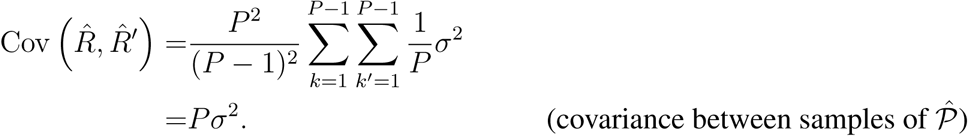

Therefore, we find that:

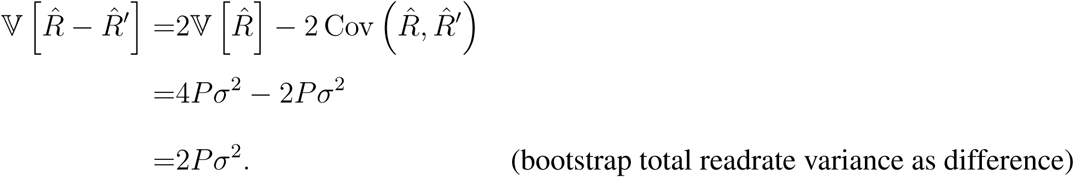

In practice, the observed nonzero positions can lead to a bootstrap distribution with variance less than its mean (underdispersed). We generally expect readrates to follow a Poisson or negative binomial (overdispersed) distribution, so in these cases, we fall back to parametric bootstrapping with a Poisson distribution with mean *µP*. Otherwise, we use the nonparametric bootstrap sampling procedure as described above.

## Supplementary Figures

**Fig. S1:**
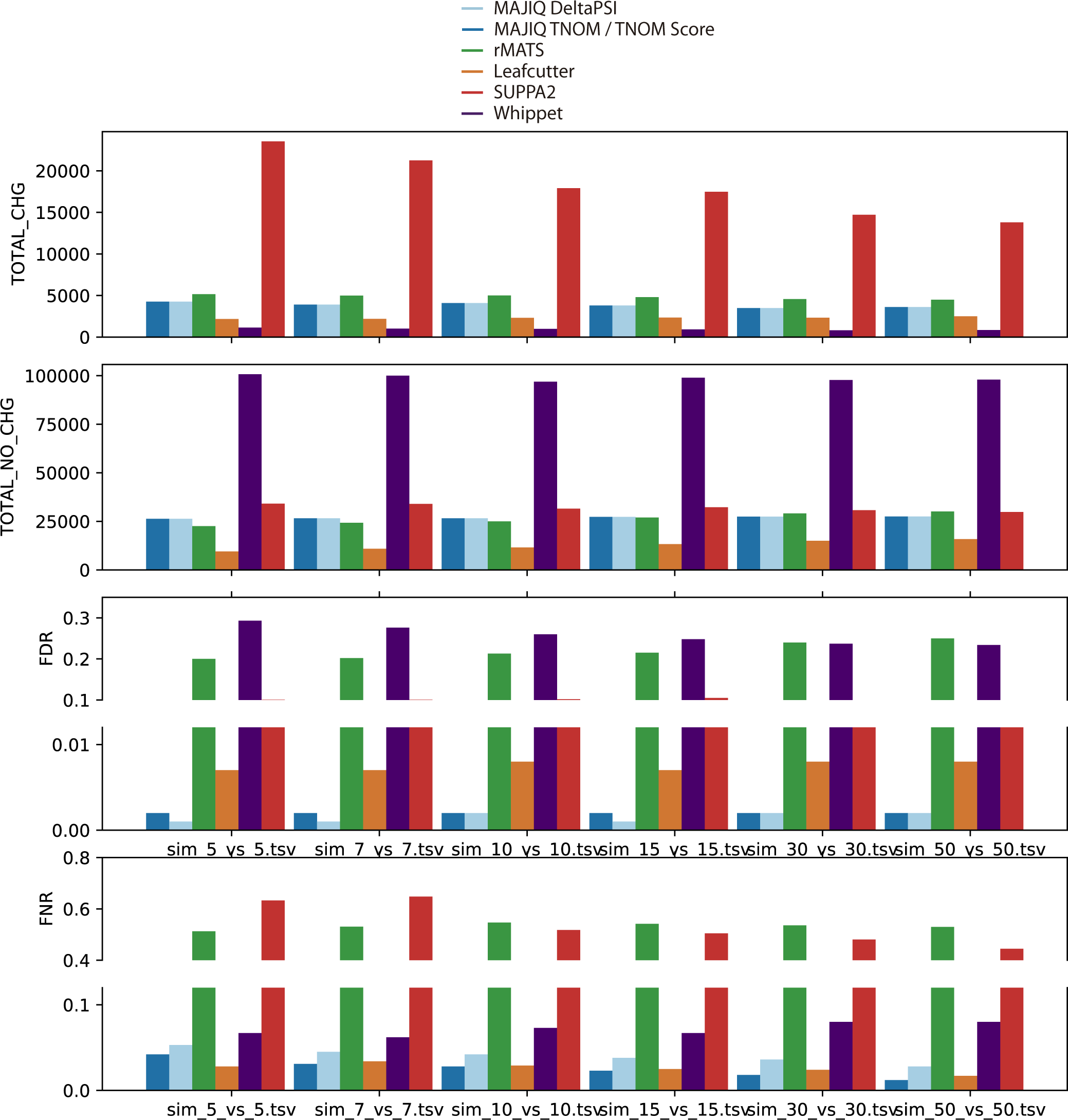
Performance evaluation using simulated data at event level. This figure is equivalent to Fig. 2B in the main text but displays the results when using each method’s unique event definition rather than aggregated at the gene level. For methods that quantify local AS events such as rMATS and MAJIQ, the number of changing events is approximately double that of changing genes (2,337 vs 4,267 for MAJIQ HET), while for LeafCutter, which uses a coarser definition for events based on intron clusters, the number of changing events and genes is similar (1,739 vs 2,169).

**Fig. S2:**
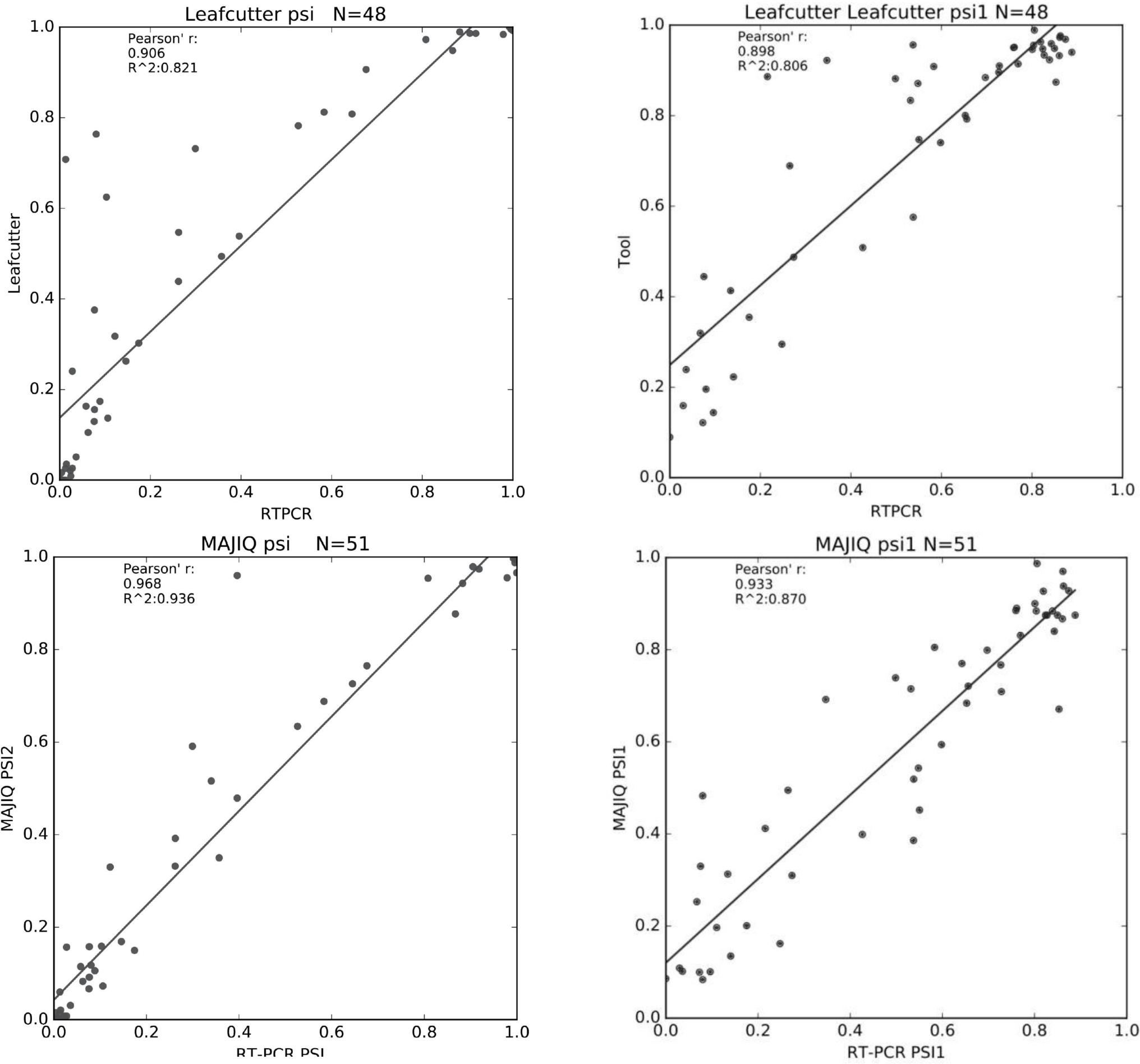
Correlation between LeafCutter and MAJIQ quantifications and RT-PCR. Correlation between RNA-seq based quantifications by LeafCutter (top row) or MAJIQ (bottom row) and RT-PCR in liver (left) and cerebellum (right). RT-PCR quantifications are from (*3*) using RNA used by (*36*) to produce the RNA-seq samples. Note that all splicing events shown here were selected by (*3*) to be binary, annotated, and changing between the two tissues to allow direct comparison to rMATS. The usage of simple binary events allowed us to calibrate LeafCutter’s intron cluster quantifications to PSI, which is not possible in the general case.

**Fig. S3:**
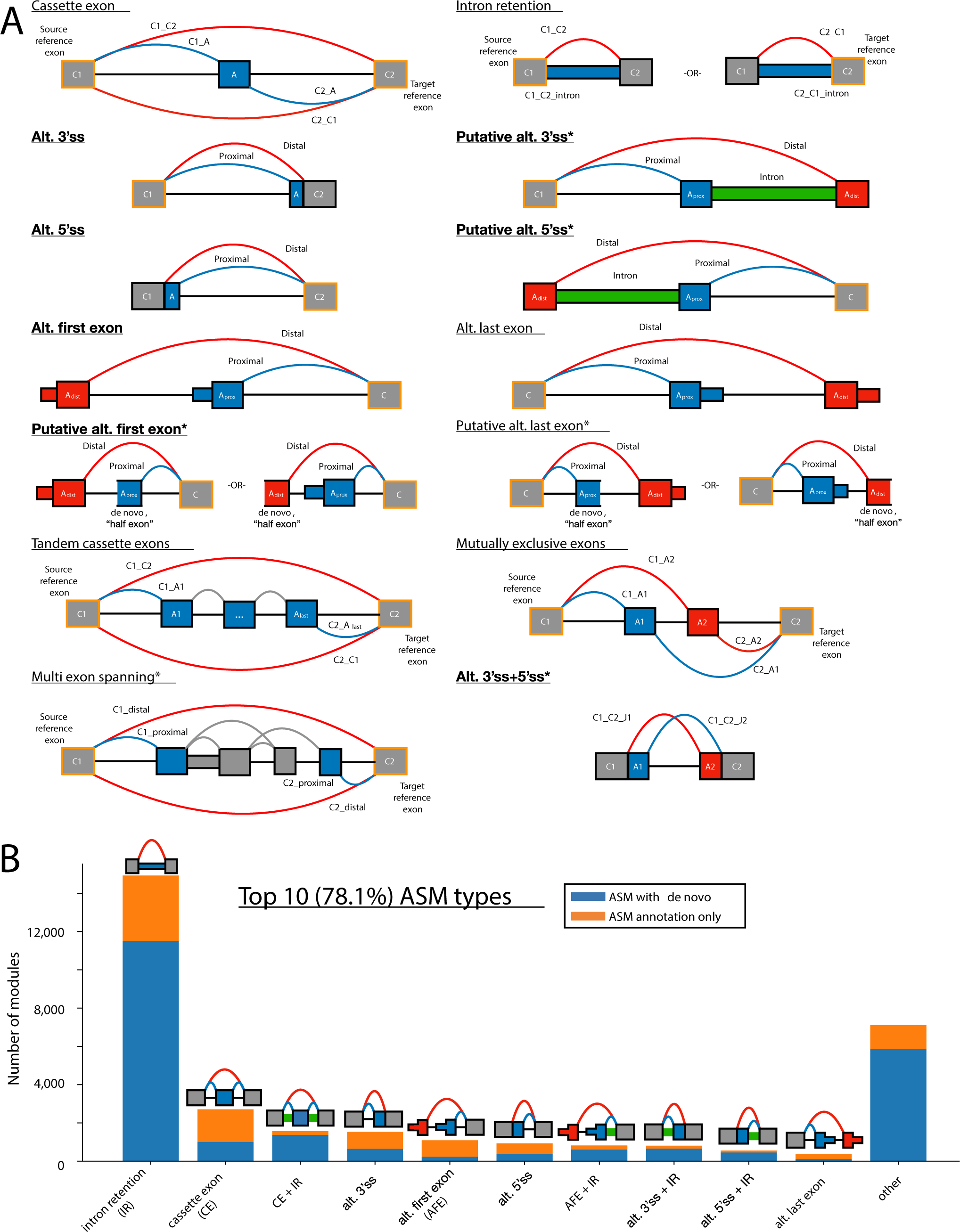
VOILA Modulizer AS event types. **(A)** Diagrams outlining the structure of alternative splicing event (AS event) types used in the VOILA Modulizer. Exons and junctions are labeled in a way consistent with the tab separated value text file outputs of the Modulizer. Grey exons outlined in orange indicate the reference exon(s) from local splicing variations (LSVs, source and/or target) used to create the splicing events. Blue junctions, introns, exons, and exonic regions correspond to inclusion products while red corresponds to exclusion products. Grey junctions in tandem cassette exons and multi-exon skipping correspond to other junctions present in the splicegraph after simplification, but are not directly considered or output by the Moduilizer in terms of quantifications. Green introns in putative 5*^′^* and 3*^′^*ss events indicate a retained intron that was quantified to high inclusion, but had the corresponding splice junction removed during simplification due to low PSI. This suggests A_prox_, the intron, and A_dist_ behave as a single exon unit with the red (intron distal) and blue (intron proximal) splice junctions acting as alternative splice sites. Asterisks indicate non-classical AS event types. **(B)** Stacked barchart showing the AS event makeup of the top 10 alternative splicing modules (ASMs) across the 13 GTEx brain tissue groups from the VOILA Modulizer after applying a 5% PSI simplification threshold (e.g. junctions with a group median of less than 5% in all groups are removed). Modules named with a plus sign (e.g. CE + IR) correspond to AS modules made up of more than one AS event type (e.g. CE + IR modules were made up of both cassette exon and intron retention events). Blue bar regions indicate AS modules that had one or more de novo or unannotated junctions, after simplification, while orange regions indicate AS modules consisting of solely annotated junctions and/or retained introns.

**Fig. S4:**
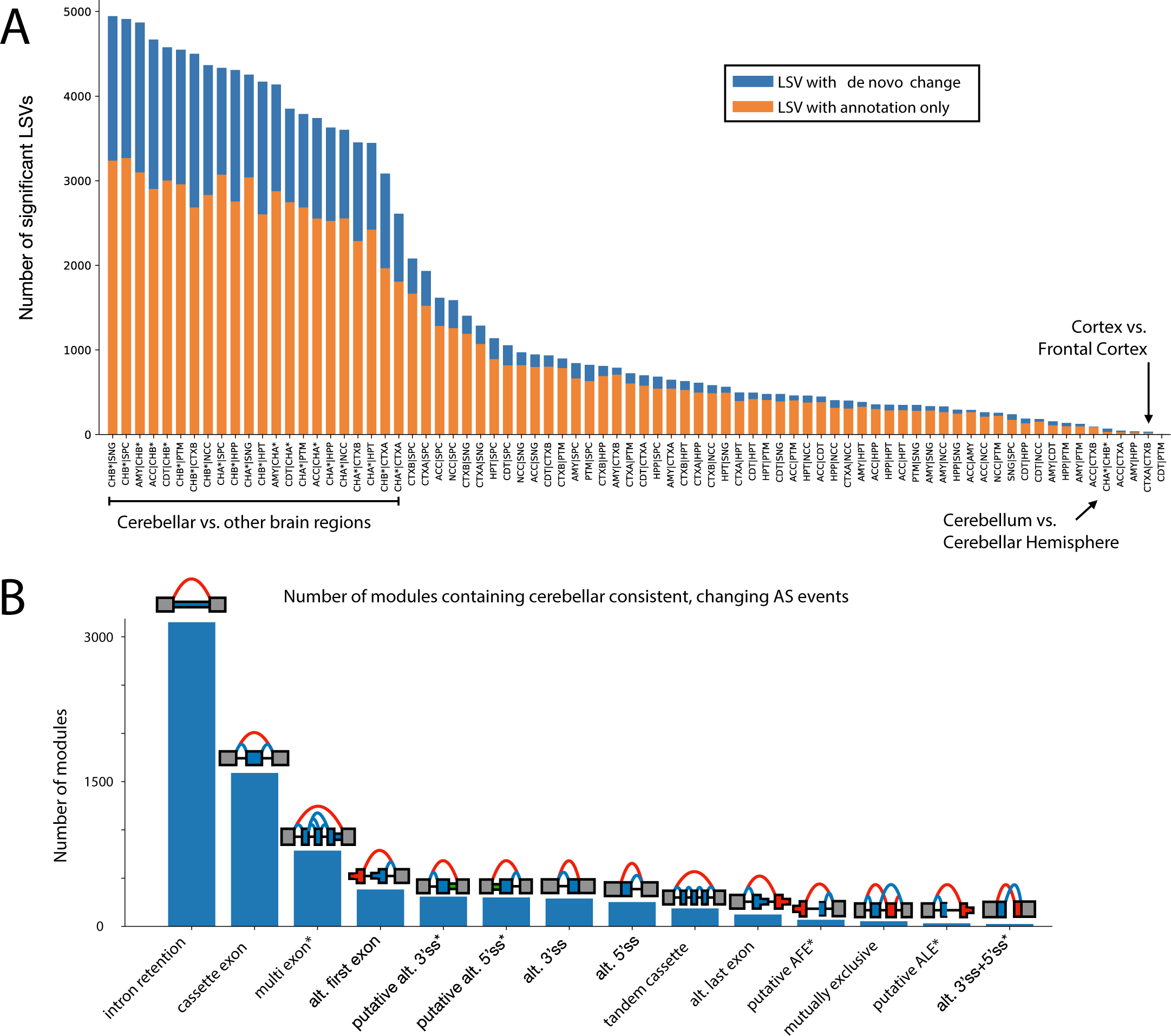
Cerebellar vs other brain tissues with LSVs and splicing modules. **(A)** Barchart showing the number of significant LSVs from 78 pairwise MAJIQ HET comparisons between the 13 GTEx brain tissue groups. Significant LSVs were those containing at least one junction or intron with an absolute difference in group median expected PSI values of 20% between two tissue groups and all four HET statistics (Wilcoxon, InfoScore, TNOM, and t-test) with *p <* 0.05. Comparisons that include a cerebellar tissue (Cerebellum, CHA; or Cerebellar Hemisphere, CHB) are highlighted. Blue indicates LSVs containing an unannotated, de novo junction/intron that was changing and orange indicates LSVs with only annotated junctions/introns. **(B)** Barchart showing the number of modules containing at least one of the 14 alternative splicing event types found to be significantly changing in both cerebellar tissues versus one or more other brain subregion tissues in a consistent way (see Methods). Event types are outlined in Figure S3A. Non-classical event types are marked with an asterisk.

**Fig. S5:**
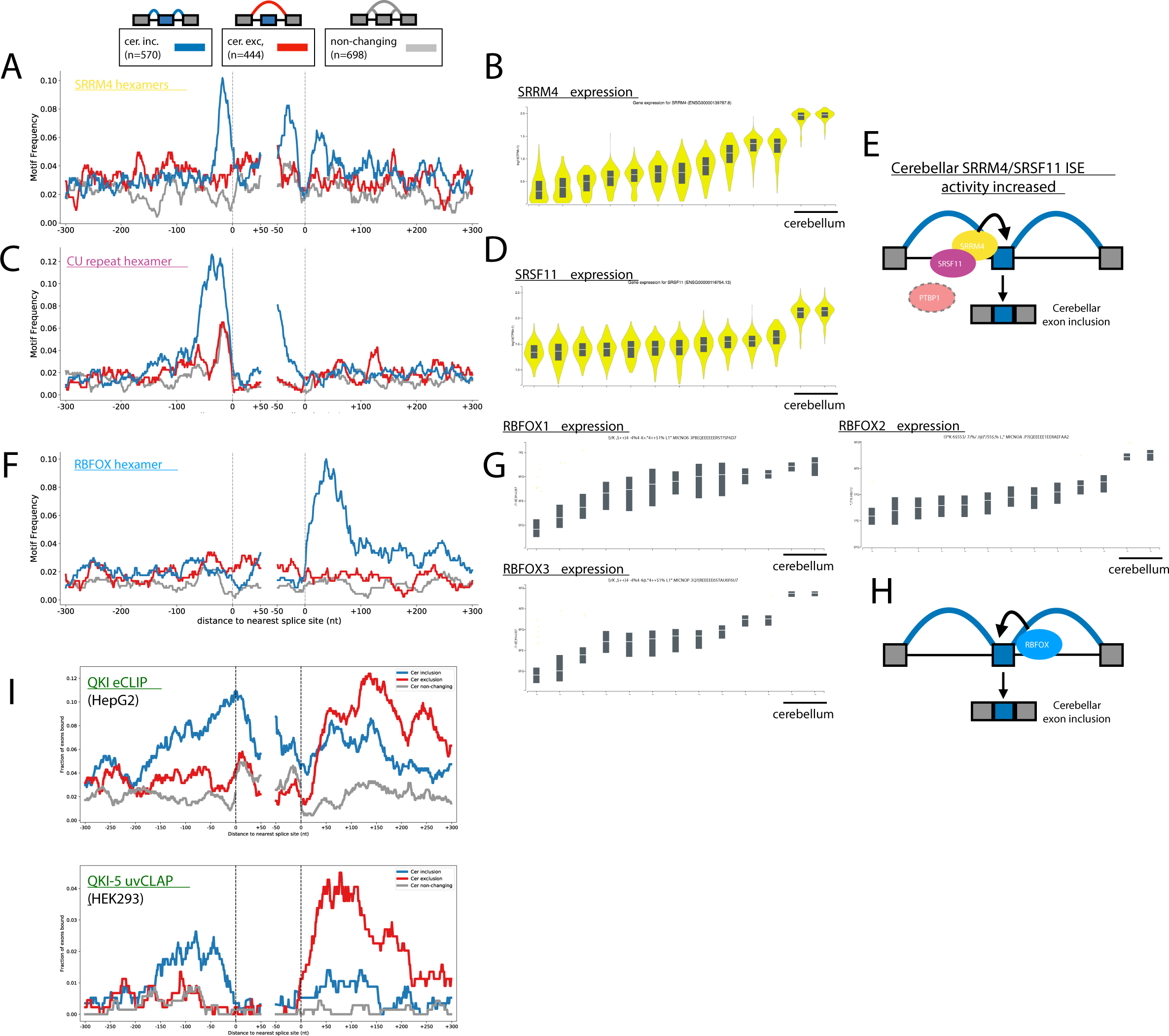
motif enrichment and RBP expression for changing and non-changing cassette exons. **(A)** RNAmaps showing the frequency of the top UGC containing SRRS6/nSR100 hexamer motifs, as determined by iCLIP (UGCUGC, CUGCUG, GCUGCC, GCUGCU (*21*)), around cerebellar inclusion (blue), exclusion (red), or non-changing (gray) CEs. Frequency was determined by searching for motif occurrence over sliding windows of 20 nucleotides with smoothing using a running mean of 5 nucleotides. **(B)** *SRRS6* bulk tissue gene expression (log_10_ (1 + TPM) for ENSG00000139767.8) sorted by median brain tissue expression. Chart generated using `GTExportal.org`. **(C)** RNAmaps showing the frequency of the top CU-repeat hexamers that bind SRSF11, as determined by iCLIP (UCUCUC and CUCUCU (*20*)), around cerebellar inclusion (blue), exclusion (red), or non-changing (gray) CEs. Frequency was determined by searching for motif occurrence over sliding windows of 20 nucleotides with smoothing using a running mean of 5 nucleotides. **(D)** *SRSF11* bulk tissue gene expression (log_10_ (1 + TPM) for ENSG00000116754.13) sorted by median brain tissue expression. Chart generated using `GTExportal.org`. **(E)** Model for SRRS6/SRSF11 promotion of exon inclusion in cerebellar tissues. Increased expression of SRRS6 and SRSF11 increases intronic splicing enhancer (ISE) activity by increased binding to CU- and UGC-rich regions just upstream of cerebellar included exons. Decreased PTB expression, which also binds CU repeat elements (*37*), may also contribute to increased SRSF11 activity. Model is based on previous work showing cooperative binding and splicing enhancement of neuronal microexons by SRSF11 and SRRS6 (*20*). **(F)** RNAmap showing the frequency of the RBFOX hexamer, UGCAUG, around cerebellar inclusion (blue), exclusion (red), or non-changing (gray) CEs. Frequency was determined by searching for motif occurrence over sliding windows of 20 nucleotides with smoothing using a running mean of 5 nucleotides. **(G)** RBFOX family bulk tissue gene expression (log_10_ (1 + TPM) for *RBFOX1*: ENSG00000078328.19, *RBFOX2*: ENSG00000100320.22, and *RBFOX3*: ENSG00000167281.18) sorted by median brain tissue expression. Chart generated using `GTExportal.org`. **(H)** Model for position dependent RBFOX regulation in GTEx brain tissues. Increased expression of RBFOX family members in cerebellar tissues leads to increased intronic splicing enhancer activity (ISE) through increased RBFOX binding downstream of exons, resulting in cerebellar exon inclusion (blue), when compared to other brain tissue groups. **(I)** RNAmaps showing the frequency of QKI CLIP peak occurrence, indicating *in vivo* binding of QKI around cerebellar inclusion (blue), exclusion (red), or non-changing (gray) CEs. Top plot shows the frequency of QKI eCLIP peaks in HepG2 cells (*34*) while bottom shows uvCLAP peak frequencies for the predominantly nuclear isoform of QKI (QK-5) in HEK293 cells that is thought to regulate splicing (*35*).

**Fig. S6:**
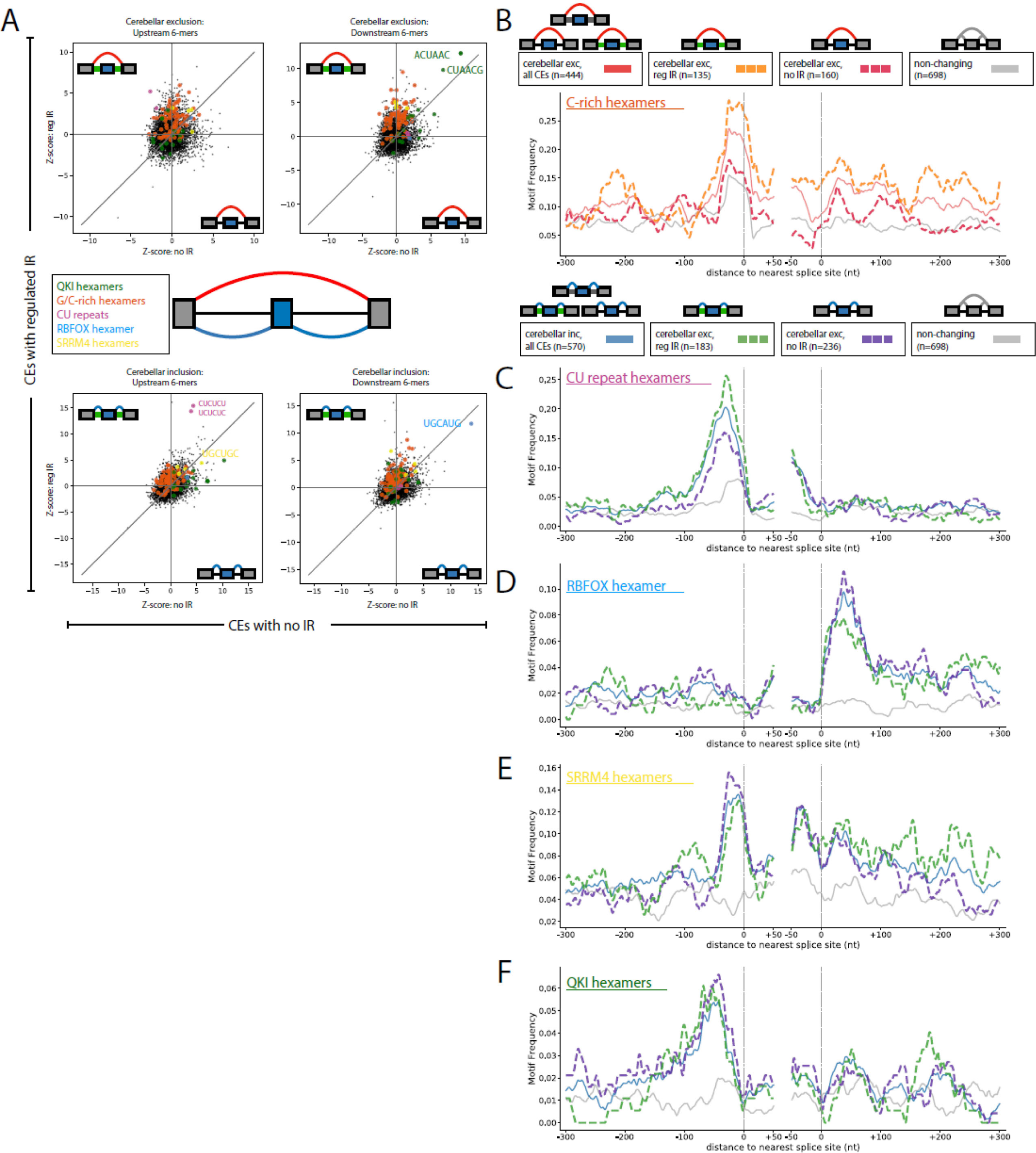
Cerebellar cassette exons with and without intron retention. **(A)** Scatter plots showing hexamer Z-score correspondence between non-overlapping sets of cerebellar cassette exon (CE) sets. Each y-axis shows Z-scores from CE events which came from AS modules containing changing intron retention (IR) event(s) versus non-changing. Each x-axis shows Z-scores from CE events coming from AS modules without IR event(s) detected. Motifs of interest are highlighted according to colors in the inset. Top plots show enrichment around cerebellar exclusion event sets while bottom plots show enrichment around cerebellar inclusion event sets. Left plots show Z-scores derived from intronic regions 300 nucleotides upstream of the 3’ss while right plots show Z-scores derived from intronic regions 300 nucleotides downstream of the 5’ss of the cassette exon. All hexamer Z-scores for various CE sets are listed in Table S1. **(B)** RNAmaps for C-rich hexamer motif for given sets of cerebellar exclusion cassette exon event sets. Lines indicate CE set according to the legend: red, all cerebellar exclusion CEs; orange dashed, subset of exclusion CEs which also contained a changing IR event; fuchsia dashed, subset of exclusion CEs with no IR event with the AS module; gray, all CEs which were not changing between comparisons. Frequency of C-rich hexamers (five of six positions are C and contain CCCC) was determined by searching for motif occurrences over sliding windows of 20 nucleotides with smoothing using a running mean of 5 nucleotides. **(C)** RNAmaps for CU-repeat hexamer motifs for given sets of cerebellar inclusion cassette exon event sets. Lines indicate CE set according to the legend: blue, all cerebellar inclusion CEs; green dashed, subset of inclusion CEs which also contained a changing IR event; purple dashed, subset of inclusion CEs with no IR event with the AS module; gray, all CEs which were not changing between comparisons. Frequency of CU-repeat hexamers (CUCUCU, UCUCUC) was determined by searching for motif occurrences over sliding windows of 20 nucleotides with smoothing using a running mean of 5 nucleotides. **(D)** Same as in (C), but shown for RBFOX hexamer (UGCAUG). **(E)** Same as in (C), but shown for SRRS6/nSR100 iCLIP hexamers (UGCUGC, CUGCUG, GCUGCC, GCUGCU (*21*)). **(F)** Same as in (C), but shown for QKI hexamers (ACUAAY).

**Fig. S7:**
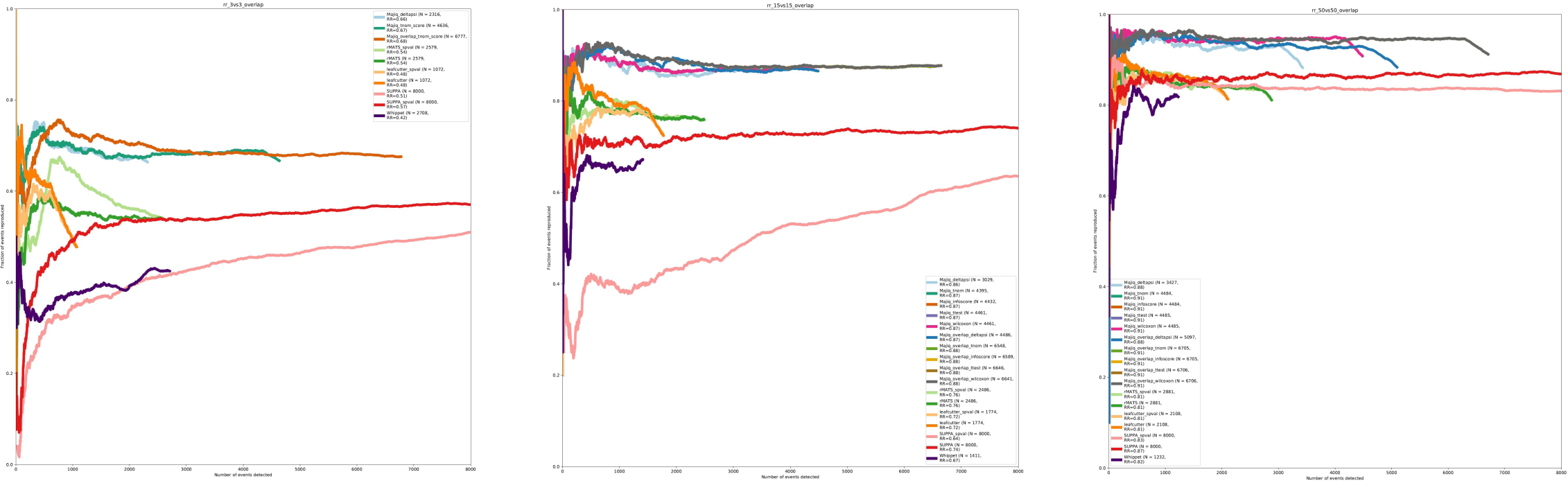
Reproducibility ratio plots without filtering MAJIQ overlapping LSVs. This figure is equivalent to Fig. 2D for reproducibility ratio plots in the main text but demonstrates the effect of not filtering MAJIQ’s list of overlapping LSVs. This filtering, used in Fig. 2D, is done to make the number of LSV events comparable to the number of classical splicing events reported by rMATS (see Methods). Note that removing the filtering step increases the number of reported differentially spliced LSVs by approximately 50% but retains similar reproducibility ratio curves.

## Supplementary Tables

The supplementary tables are provided as a Microsoft Excel workbook with sheets prefixed by “Table” and “README”. The “Table” sheets specify the tables themselves, and for each table, there is a corresponding “README” sheet which describes the format/columns of the table.

**Table S1: Hexamer Z scores for cerebellar CE sets** Z-scores calculated for hexamer occurence within 300 nucleotides upstream or downstream of cerebellar cassette exon sets versus a set of stringent non-changing cassette exons.

**Table S2: Tool parameters for performance evaluations** The tools, versions, and additional parameters used for the performance evaluations.

## Notes

### Summary of Updates

Fixed minors typos and updated format.

